# Inhibition of TRPV4 rescues circuit and social deficits unmasked by acute inflammatory response in a Shank3 mouse model of Autism

**DOI:** 10.1101/2021.10.13.464215

**Authors:** Stamatina Tzanoulinou, Stefano Musardo, Alessandro Contestabile, Sebastiano Bariselli, Giulia Casarotto, Elia Magrinelli, Yong-hui Jiang, Denis Jabaudon, Camilla Bellone

**Author notes:** Equal contribution (Stamatina Tzanoulinou, Stefano Musardo, Alessandro Contestabile). Department of Biomedical Sciences (DSB), FBM, University of Lausanne, Lausanne, Switzerland.

## Abstract

Autism spectrum disorder is a neurodevelopmental disease characterized by social deficits and repetitive behaviors. The high heterogeneity of the disease may be explained by gene and environmental interactions and potential risk factors include immune dysfunctions and immune-mediated co-morbidities. Mutations in the *SHANK3* gene have been recognized as a genetic risk factor for ASD. While heterozygous *SHANK3* mutations are usually the types of mutations associated with idiopathic autism in patients, heterozygous deletion of *Shank3* gene in mice does not commonly induce ASD-related behavioural deficit. Here, we used *in-vivo* and *ex-vivo* approaches to demonstrate that region-specific neonatal downregulation of *Shank3* in the NAc promotes D1R-MSN hyperexcitability and upregulates *Trpv4* to impair social behaviour. Interestingly, genetically vulnerable *Shank3^+/-^* mice, when challenged with Lipopolysaccharide to induce inflammatory response, showed similar circuit and behavioural alterations that were rescued by acute Trpv4 inhibition. Altogether our data demonstrate shared molecular and circuit mechanisms between ASD-relevant genetic alterations and environmental insults, which ultimately lead to sociability dysfunctions.

## Introduction

Autism spectrum disorder (ASD) includes a heterogeneous group of neurodevelopmental diseases characterized by social communication deficits and repetitive behaviours. Mutations in *SHANK3* gene, coding for a scaffolding protein located at excitatory synapses, account for 1-2% of all ASD cases and its haplo-insufficiency is acknowledged to lead to a high-penetrance form of ASD, known as Phelan-McDermid syndrome^1, 2^. Currently, the development of pharmacological interventions to alleviate ASD-related sociability symptoms is limited by several factors, including the relative lack of understanding of the genetic consequences of *SHANK3* insufficiency. This is further complicated by the fact that Shank3 plays specific roles depending on its expression pattern in different regions and cell types^3^. Thus, investigating altered neuronal circuit mechanisms underlying disease pathophysiology and uncovering their roles in discrete behavioral readouts in mice is of the highest importance^4, 5^.

Most of the pre-clinical models of *Shank3* deficiency show impairments in dorsal striatal circuits^3, 6^, principally related to the indirect pathway, which drives repetitive behaviour^6, 7^. On the other hand, the role of the mesolimbic reward system, including the Ventral Tegmental Area^8^ and the Nucleus Accumbens (NAc)^9^, in social reward processing makes it an ideal neural circuit substrate for further investigation in the context of ASD in both rodents^10^ and humans^11, 12^. Despite the fact that neuronal deficits within the reward system have been revealed in different *Shank3* animal models^13, 14^ and that expression of Shank3 in the striatum is particularly enriched^15^, the contribution of *Shank3* insufficiency in the ventral striatum, which includes the Nucleus Accumbens (NAc), to ASD symptomatology has been largely neglected.

Although the generation of knock-out (KO) *Shank3* (*Shank3^-/-^*) mouse lines has favored the identification of behavioural and synaptic impairments, single allele mutations minimally affect the behavioural pattern in rodents^16–19^ limiting translation from rodents to human studies. Indeed, while Phelan-McDermid syndrome (PMS) patients are heterozygous for *SHANK3* deletions or mutations, most of the existing animal models failed to report consistent behavioural phenotypes when heterozygote mice were assessed. Thus, one intriguing question that arises is whether environmental challenges would actually exacerbate or unmask alterations, otherwise covert in heterozygote mice. Indeed, apart from genetic risk factors, several studies support the role of immune regulation and inflammation in ASD. Patients have frequent immune dysfunctions and immune-mediated co-morbidities^20^. Furthermore, transcriptomic analysis in post-mortem brain tissues revealed an upregulation of genes involved in inflammation ^21^, while in recent years, an interplay between immune system and reward circuit function has been put forward^22, 23^. Remarkably, in some PMS patients, debilitating symptoms appeared after acute infections or stressful environmental challenges ^24^ suggesting that the heterogeneity in the phenotypes could be the consequence of the interplay between genetic and environmental factors. Although increasing evidence indicates links between immune deficits and ASD, mechanistic insights are still lacking.

Here we firstly interrogated behavioural and electrophysiological consequences of shRNA-induced *Shank3* early postnatal downregulation in the NAc. Not only we observed reduced social preference and D1-medium spiny neurons (D1-MSNs) hyperexcitability, but also identified the Transient Receptor Potential Vanilloid 4 (Trpv4) as a key effector of our observations. Remarkably, similar molecular, circuit and behavioural alterations were also observed in genetically vulnerable *Shank3*^+/-^ mice challenged with lipopolysaccharide-induced neuroinflammation. Finally, acute Trpv4 inhibition in the NAc restored excitability and sociability deficits in *Shank3* heterozygous mice.

## RESULTS

### Social deficits following early NAc-specific *Shank3* insufficiency

Given the emerging importance of NAc in social reward processing^9, 25^, we first focused our investigation on this brain circuit and asked whether *Shank3* downregulation restricted to this region would lead to sociability deficits. Using AAV-sh*Shank3*-luczsGreen virus, we downregulated the expression of *Shank3* during early postnatal development^13, 26^ (≤P6; hereafter P6 for simplicity, **Fig. 1a-a’ and Supplementary figure 1a-b**) or during adulthood (P90, **Fig. 1d-d’, and Supplementary figure 1a-b**). While scr*Shank3* mice (injected at the same age with a scrambled virus) showed preference for a juvenile conspecific (**Fig. 1b** and **1e)**, sh*Shank3* P6-injected mice spent a comparable amount of time around the juvenile and object stimulus (**Fig. 1c)** and in the two chambers (**Supplementary figure 1c**). Furthermore, while the overall exploratory behaviour was comparable between scr*Shank3* and sh*Shank3* mice **(Supplementary figure 1d and g**), sh*Shank3*-infected mice spent less time exploring the juvenile mouse and more time exploring the object stimulus compared to scr*Shank3* (**Supplementary figure 1e-f**). Remarkably, when shShank3 was injected at P90, mice showed intact sociability **(Fig. 1f and Supplementary figure 1h**) while presenting similar decrease in Shank3 expression (**Supplementary figure 1b**). No difference in the exploration time around targets between groups (**Supplementary figure 1i-k**), nor in the distance moved **(Supplementary figure 1l)** was observed after injection at P90.

**Figure 1:**
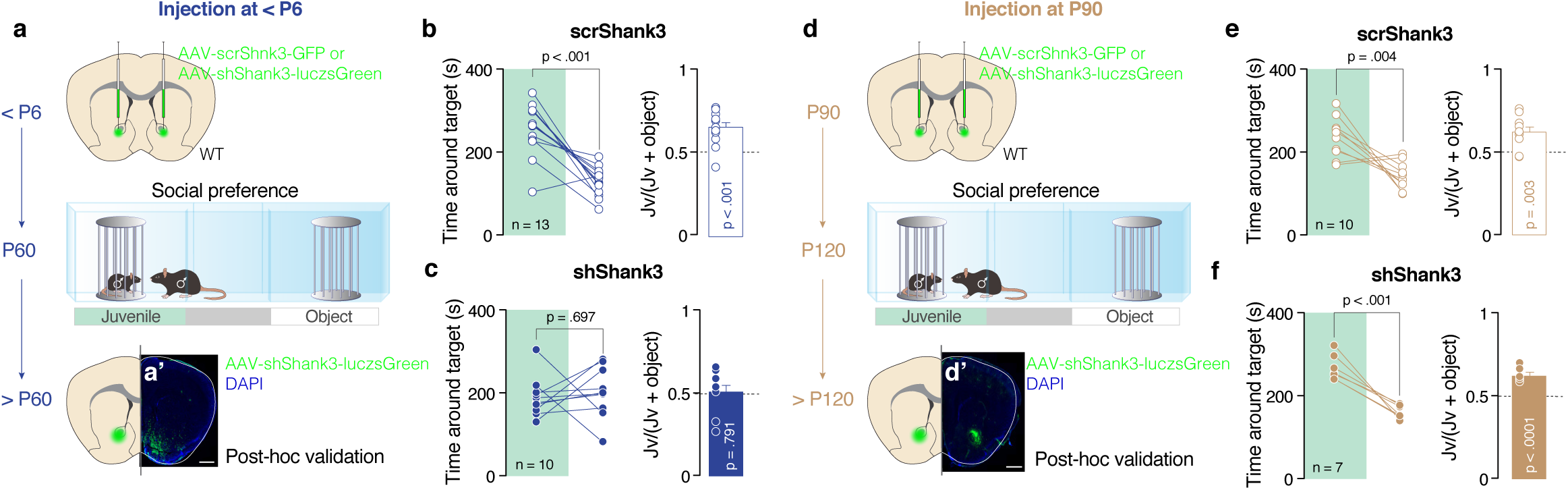
*D*ownregulation of *Shank3* in the NAc during early postnatal development alters social preference. **(a, d)** Schema of injection sites in the NAc with AAV-scr*Shank3*-GFP or AAV-sh*Shank3*-luczsGreen in ≤P6 mice (e) or at P90 (h). **(a’, d’)** Representative images of injection sites (scale bar: 500 µm). **(b, c, e, f)** Left: time spent around the enclosures during the social preference test for mice injected at ≤P6 or at P90 ((b); *t* _(12)_ = 6.092, *p* < 0.001, (c); *t* _(9)_ = 0.409, *p* = 0.697, (e); *t* _(9)_ = 3.806, *p* = 0.004, (f); *t* _(6)_ = 6.970, *p* < 0.001). Right: juvenile preference index for mice injected at ≤P6 or at P90 (one-sample t-tests against chance level = 0.5: (b), scr*Shank3*; *t* _(12)_ = 5.847, *p* < 0.001, (c), sh*Shank3*; *t* _(9)_ = 0.273, *p* = 0.791; (e), scr*Shank3*; *t* _(9)_ = 3.928, *p* = 0.003, (f), shShank3; *t* _(6)_ = 7.996, *p* < 0.001). Error bars report SEM.

Altogether these data point at the NAc as a key region for sociability deficits induced by *Shank3* insufficiency. Moreover, our results suggest the existence of a critical period during early postnatal development, which is important for the expression of appropriate sociability later in life. We, thus, decided to focus our efforts on early *Shank3* downregulation and to investigate the mechanisms underlying sociability deficits.

### Alterations in intrinsic properties of NAc D1R-expressing Medium Spiny Neurons following *Shank3* downregulation

The downregulation of Shank3 from D1R-expressing (direct pathway) or D2R-expressing MSNs of dorsal striatum (indirect pathway) leads to neuronal hyperexcitability^3^. In order to assess the excitability of MSN subpopulations in the NAc in our model, we used fluorescently-labelled D1R mice (Drd1a-tdTomato)^27^ injected with scr*Shank3* or sh*Shank3* (**Fig. 2a-a’**). When recorded in presence of synaptic blockers (picrotoxin and kynurenic acid), *Shank3* downregulation increased the excitability of D1R-tom^+^ compared to scr*Shank3*::D1R-tom^+^ MSNs, while no changes were detected in the D1R-tom^-^population (**Fig. 2b-e** and **Supplementary figure 2a-f**). Interestingly, in absence of synaptic blockers, *Shank3* downregulation induced a hyperexcitability of D1R-tom^+^ MSNs and hypoexcitability of D1R-tom^-^ MSNs, (**Supplementary figure 2g-r**). Overall, these results indicate that the hyperexcitability of accumbal direct pathway MSNs largely derives from alterations of intrinsic membrane properties, while the hypoexcitability of putative indirect pathway MSNs is the consequence of circuit network dysfunctions.

**Figure 2:**
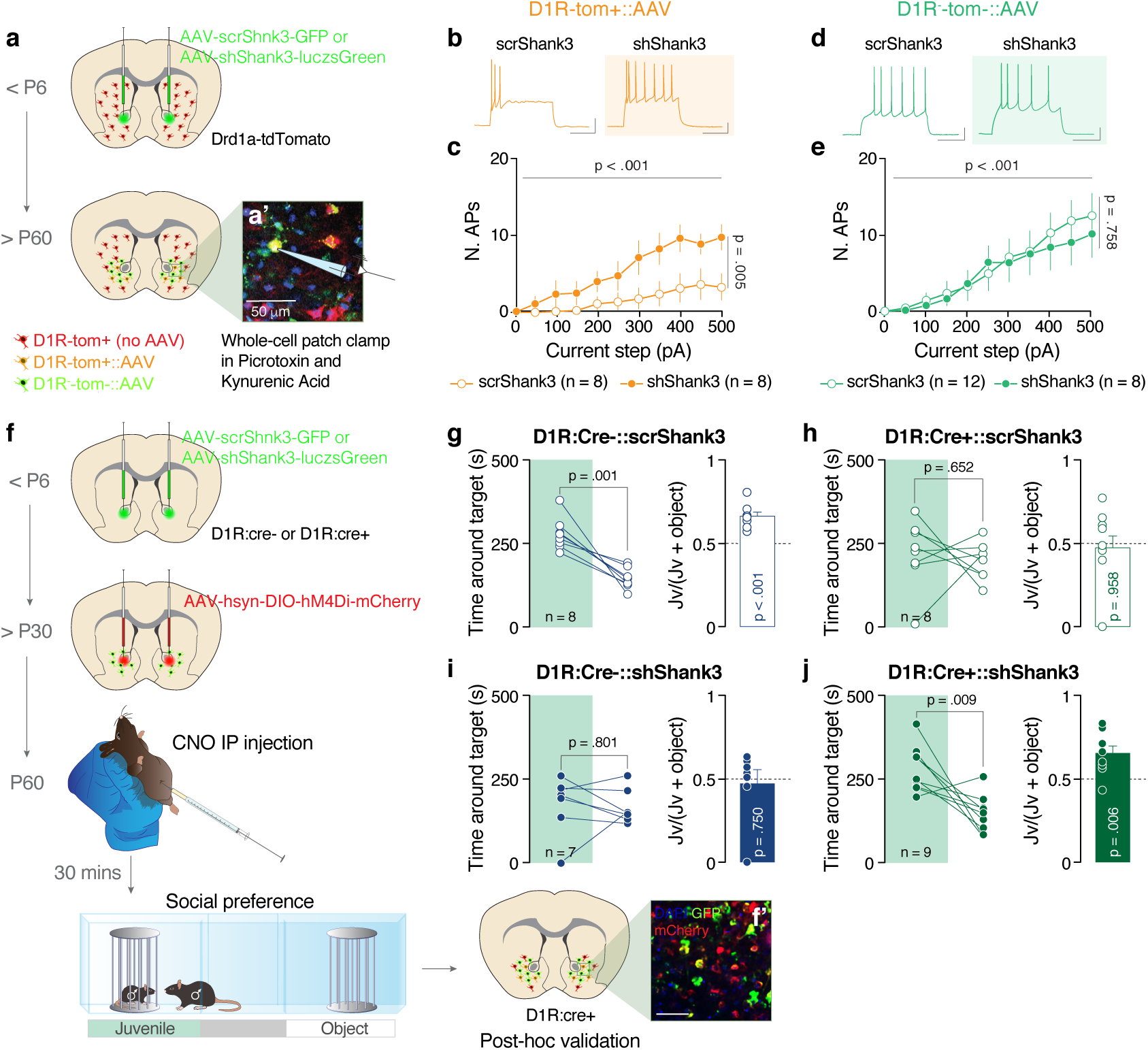
*Shank3* NAc downregulation alters D1R MSNs excitability. Decreasing the activity of D1R MSNs normalizes sociability deficits. **(a)** Experimental design. Drd1a-dTomato mice were injected neonatally in the NAc with scr or sh*Shank3* virus and whole-cell patch clamp recordings were performed during early adulthood. **(a’)** Representative images of the NAc of a D1R-tom+::sh*Shank3* mouse (scale bar: 50 µm). **(b)** Example traces at 300 pA depolarizing current injection in D1R-tom+ MSNs infected with scr*Shank3* (left) or with sh*Shank3* (right). **(c)** Number of action potentials (nAPs) across increasing depolarizing current steps (0-500 pA) for D1R-tom+::scr*Shank3* and sh*Shank3* MSNs, in presence of picrotoxin and kynurenic acid. (Repeated measures ANOVA, virus main effect *F _(1,14)_ = 10.88, p =* 0.005, current steps main effect *F _(10, 140)_ = 7.727, p <* 0.001, scr*Shank3* n = 8 cells, 3 mice, sh*Shank3* n = 8 cells, 3 mice). **(d)** Example traces at 300 pA depolarizing current injection in D1R-tom-MSNs infected with scr*Shank3* (left) or with sh*Shank3* (right). **(e)** Number of action potential (nAPs) across increasing depolarizing current steps (0-500 pA) for D1R-tom-::scr*Shank3* and sh*Shank3* MSNs, in presence of picrotoxin and kynurenic acid. (Repeated measure (RM) two-way ANOVA, virus main effect *F _(1,18)_ =* 0.098, *p =* 0.758, current steps main effect *F _(10 , 180)_ =* 14.58, *p <* 0.001, n = 8 cells, 3 mice (sh), n=12 cells, 3 mice (scr)). **(f)** Experimental design. D1R-Cre positive (D1R:Cre^+^) and negative (D1R:Cre^-^) mice were injected neonatally in the NAc with scr or sh*Shank3* virus and after P30 with AAV-hSyn-DIO-hM4Di-mCherry (DREADD). After 4 weeks, allowing for virus expression, the mice underwent social behaviour assessment in the three-chamber task. Mice were intraperitoneally injected with CNO 30 min before starting the test. **(f’)** Representative images of the NAc of a D1R:Cre^+^ mouse infected with sh*Shank3* and DREADD viruses (scale bar: 50 µm). **(g, h, i, j**) Left: time around the target during the social preference test for D1R:Cre^-^::scr*Shank3* mice (g; *t* _(7)_ = 5.453, *p* = 0.001); D1R:Cre^+^::scr*Shank3* mice (h; *t* _(7)_ = 0.471, *p* = 0.652); D1R:Cre^-^ ::sh*Shank3* mice (i; *t* _(6)_ = 0.264, *p* = 0.801) and D1R:Cre^+^::sh*Shank3* mice (j; *t* _(8)_ = 3.443, *p* = 0.009). Right: juvenile preference index (one-sample t-tests against chance level = 0.5: D1R:Cre^-^::scr*Shank3*; *t* _(7)_ = 6.395, *p* < 0.001, D1R:Cre^+^::scr*Shank3*; *t* _(7)_ = 0.054, *p* = 0.958, D1R:Cre^-^::sh*Shank3*; *t* _(6)_ = 0.334, *p* = 0.750, D1R:Cre^+^::sh*Shank3*; *t* _(8)_ = 3.706, *p* = 0.006). Error bars report SEM.

### A causal link between NAc D1R MSN hyperexcitability and sociability deficits

To probe causality between direct pathway hyperexcitability and sociability deficits, we used cell-specific chemogenetic tools to manipulate neuronal activity. D1R:Cre^+^ or D1R:Cre^-^ mice were injected with either control scrambled virus or sh*Shank3* during early post-natal development. After P30, all mice were infected with an inhibitory Cre-dependent DREADD-expressing virus (AAV-DIO-hM4Di-mCherry) (**Fig. 2f-f’**). We validated the effectiveness of our chemogenetic approach by analyzing the expression of GIRK channels in Nac MSN (**Supplementary figure 3a**) the main effectors of chemogenetic inhibition, and the effects of Clozapine N-Oxide (CNO) on neuronal excitability *ex vivo* (**Supplementary figure 3b-c**). Mice underwent the three-chamber interaction assay 30 minutes after systemic CNO injections (**Fig. 2f-f’**). D1R:Cre^+^ mice injected with CNO (regardless of NAc virus) reduced their locomotor activity (**Supplementary figure 3d**); however, the total exploration time for both enclosures remained comparable to that of D1R:Cre^-^ mice (**Supplementary figure 3e**). By analysing the time spent around either the juvenile or object target, we confirmed that control D1R:Cre^-^::scr*Shank*3 mice showed intact sociability (**Fig. 2g, Supplementary figure 3f**). While both D1R:Cre^+^::scr*Shank*3 mice treated with CNO and D1R:Cre^-^::sh*Shank*3 did not show a preference for the social over the object stimulus (**Fig. 2h-i and Supplementary figure 3f**), interestingly, D1R:Cre^+^::shShank3 showed a preference for the social stimulus compared to the object (**Fig. 2j and Supplementary figure 3f**).

These data establish a causal link between NAc D1R-MSN hyperexcitability and sociability defects *in vivo*, and further suggest that decreasing direct pathway hyperexcitability might be a useful strategy to ameliorate social dysfunctions.

### Downregulation of *Shank3* induces a *Trpv4* upregulation

It has been previously shown that Shank3 mutations in human predispose to autism by inducing a channelopathy^28^. To further investigate the mechanisms underlying D1-MSN hyperexcitability, we performed direct pathway transcriptomic analysis of the NAc in Drd1a-tdTomato P6-injected scr*Shank*3 and sh*Shank*3 mice. For this purpose, we FAC-sorted direct pathway MSNs at P30 and performed bulk RNA sequencing (**Fig. 3a**). Scr*Shank3* and sh*Shank3* were clustered separately in both D1R-tom^+^ and D1R-tom^-^ populations, (**Fig. 3b**) and differential expression analysis by groups for scr*Shank3 vs* sh*Shank3* revealed 178 altered genes in AAV-infected D1R-tom^+^ (**Fig. 3c and Supplementary figure 4a**). GO:Term analysis of significantly altered genes in NAc-injected shShank3 mice, highlighted changes relevant to cell adhesion, localization, and cellular movement-related, as well as, related to functions regarding inflammatory mechanisms (**Fig. 3d**). Moreover, within the modified genes identified in the bulk RNA sequencing, we observed a high representation of activity-related genes (**Fig. 3e**) in D1R-tom^+^ neurons, immune response-related genes, as well as, SFARI genes associated with ASD in both D1R-tom^+^ and D1R-tom^-^ populations (**Supplementary figure 4b-c**).

**Figure 3:**
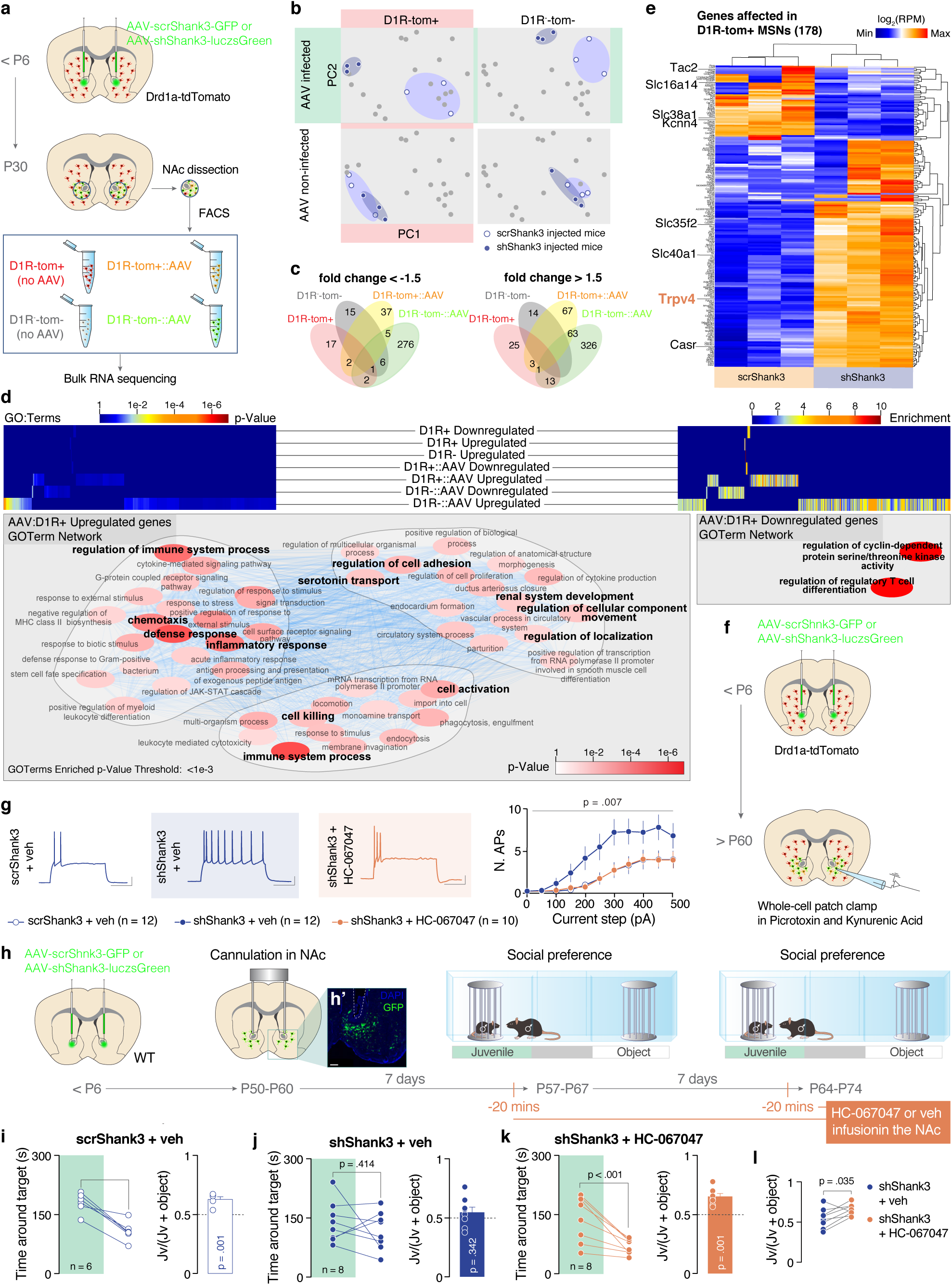
Downregulation of *Shank3* in D1R MSNs induces alterations in inflammatory mediators and Trpv4 expression. **(a)** Experimental design. Drd1a-dTomato mice were injected neonatally in the NAc with scr or sh*Shank3* virus. At P30 NAc was dissected and FACsorted in 4 different cell populations (D1R-tom+, D1R-tom-, D1R-tom+::AAV and D1R-tom-::AAV). For each cell population we carried out bulk RNA sequencing. **(b)** Worst-case scenario selected altered genes in scr vs sh testing clearly discriminated infected cells, both D1R+ and D1R- in PCA analysis. **(c)** While non-infected samples do not share common genes significantly altered in scr vs sh testing, infected D1R+ and D1R- share a core set of 68 altered genes. **(d)** Overall GO:Term analysis of infected D1R+ significantly altered genes highlights the relevance of inflammatory mechanisms, as well as cell adhesion-, localization- and movement-related functions. **(e)** D1R-tom+ altered genes include genes expressing proteins directly involved in electrophysiological properties, including the Transient receptor potential vanilloid 4 (*Trpv4*). **(f)** Experimental design. Drd1a-dTomato mice were injected neonatally in the NAc with scr or sh*Shank3* virus and whole-cell patch clamp recordings were performed during early adulthood. **(g)** Right: example traces from 300 pA depolarizing current injection in D1R+ MSNs infected with scr*Shank3* treated with vehicle (up), D1R+ MSNs infected with sh*Shank3* treated with vehicle (middle) and D1R+ MSNs infected with sh*Shank3* treated with HC-067047 (down). Left: number of action potentials (nAPs) across increasing depolarizing current steps (0-500 pA) for D1R-tom+::scr*Shank3* and sh*Shank3* MSNs in the presence of Trpv4 antagonist (HC-067047). (Repeated measures ANOVA, drug main effect *F _(2, 31)_ =* 5.883, *p* = 0.007, current steps main effect *F _(10 , 310)_ =* 24.15, *p <* 0.001, drug by current steps interaction *F _(20 , 310)_ =* 1.685, *p =* 0.035 n = 12 cells, 4 mice (sh*Shank3*-Veh), n=10 cells, 3 mice (sh*Shank3*-Trpv4), n = 12 cells, 4 mice (scr*Shank3*-Veh)). **(h)** Experimental design. C57BL6/j mice were injected neonatally in the NAc with scr or sh*Shank3* virus and at P50-60 were bilaterally cannulated above the NAc. After 7 days, mice underwent the three-chamber social interaction assay. Scr*Shank3* were infused with vehicle (aCSF/DMSO 0.3%). On the other hand, sh*Shank3* mice were infused with either vehicle (aCSF/DMSO 0.3%) or HC-067047 (2µg in aCSF/DMSO 0.3%) 10 min before to start the test. **(h’)** Representative image of the injection site and cannula placement above the NAc (scale bar: 250 µm). **(i, j, k)** Left: time around the target during the social preference test for mice infected with scr*Shank3* and infused with vehicle (i, *t* _(5)_ = 6.304, *p* = 0.002), mice infected with sh*Shank3* and infused with vehicle (j, *t* _(7)_ = 0.869, *p* = 0.414) or with HC-067047 (k, *t* _(7)_ = 4.324, *p* = 0.004). Right: juvenile preference index for mice infused either with vehicle or with HC-067047 (one-sample t-tests against chance level = 0.5: (f); *t* _(5)_ = 6.459, *p* = 0.001, (g); *t* _(7)_ = 1.02, *p* = 0.342, (h); *t* _(7)_ = 6.078, *p* = 0.001). **(l)** Juvenile preference index comparison between sh*Shank3*-vehicle and sh*Shank3*-HC-067047 (*t* _(7)_ = 2.6, *p* = 0.035). Error bars report SEM.

Among these genes altered by early postnatal *Shank3* downregulation, we noticed that the one encoding for the Transient receptor potential vanilloid 4 (*Trpv4*) channel was significantly upregulated (**Fig. 3e)**. Trpv4 is a member of the transient receptor potential superfamily, broadly expressed in the central nervous system^29^. These receptors are activated by temperature, mechanical stimulation, cell swelling and endocannabinoids^30^ and participate in inflammatory responses^31^. Moreover, Trpv4 function influences neuronal excitability and its disruption leads to social behaviour abnormalities^32^. The increased in Trpv4 expression in the NAc from mice where Shank3 was downregulated before P6, was confirmed by qPCR (**Supplementary figure 5a)**. Remarkably, when Shank3 was downregulated during adulthood, the levels of Trpv4 were comparable between scrShank3 and shShank3 injected mice (**Supplementary figure 5b**). Since downregulation of *Shank3* in adulthood did not reveal any sociability deficit (**Fig. 1d-f**), together, these data suggested a link between the increased *Trpv4* expression and the behavioural phenotype. To directly interrogate this hypothesis, we tested the ability of a Trpv4-specific inhibitor (HC-067047) to rescue the direct pathway MSN hyperexcitability *ex-vivo* (**Fig. 3f**). In patch-clamp recordings, bath application of HC-067047 normalized the excitability of D1R-tom^+^::sh*Shank3* to D1R-tom^+^::scr*Shank3* levels (**Fig. 3g**). So far, this evidence indicates that early downregulation of *Shank3* in the NAc upregulates both the expression and the function of *Trpv4* in the direct pathway neurons, identifying a novel molecular effector of *Shank3* insufficiency.

### Trpv4 antagonist restores sociability in NAc sh*Shank3* mice

To test causality between sociability defects and the upregulation of *Trpv4* in the NAc and to probe its potential as a therapeutic target *in-vivo*, we next asked whether the region-specific administration of Trpv4 inhibitor restores sociability in sh*Shank3* mice. Scr*Shank3* and sh*Shank3* were bilaterally cannulated above the NAc for local pharmacology experiments. After one week of recovery, sh*Shank3* mice were pre-treated with HC-067047 or vehicle before the three-chamber test (**Fig. 3h-h’**). Treatments were counterbalanced and the same animals were tested again after seven days (scr*Shank3* animals were instead infused only with vehicle). scr*Shank3* mice infused with vehicle showed intact sociability (**Fig. 3i**) indicating no side effects of the cannulation on our behavioural endpoints. Confirming our previous findings, vehicle-infused sh*Shank3* animals showed impaired social preference (**Fig. 3j**). Remarkably, intra-NAc Trpv4 antagonist (HC-067047) infusions in sh*Shank3* mice restored sociability (**Fig. 3k**), increasing the time spent in the social chamber (**Supplementary figure 5c**). Furthermore, sh*Shank3* mice showed an increase of social preference ratio when infused with HC-067047 compared to when infused with vehicle (**Supplementary figure 3l**). No difference was observed in the distance moved during the test among the groups (**Supplementary figure 5d**).

Our results highlight the role of Trpv4 both in D1R-MSN hyperexcitability and social preference deficits displayed by NAc-injected sh*Shank3* mice.

### LPS challenge unmasks social deficits in *Shank3^+/-^* mice

Based on the region-specific results obtained by GO:Term analysis (**Fig. 3d**), we hypothesised that acute inflammatory challenge could unmask behavioural deficits of *Shank3* knock-out mice, in which exons 4 to 22 were deleted (Δe4-22^+/-^ mice, hereafter referred to as Shank3^+/-^)^19^, via a Trpv4-dependent mechanism. As previously reported^19^, *Shank3^+/-^* mice do not display social preference deficits in the three-chamber test (**Supplementary figure 6a-e**). Importantly, when Lipopolysaccharide (LPS) was injected 24 hrs prior the three-chamber task (**Fig. 4a**), *Shank3^+/-^* mice spent a comparable amount of time around the juvenile and object stimulus and in the corresponding chamber, indicating sociability deficits (**Fig. 4e** and **Supplementary figure 7a**). As control, saline-injected *Shank3^+/+^* and *Shank3^+/-^* mice spent more time exploring the juvenile-containing enclosure and chamber (**Fig. 4b** and **d** and **Supplementary figure 7a**). Moreover, LPS challenge did not confer any behavioural alterations in *Shank3^+/+^* mice (**Fig. 4c** and **Supplementary figure 7a**) and the distance moved did not differ across genotypes (**Supplementary figure 7b**). Importantly, sociability deficits were not observed 7 days after LPS injection (**Fig. 4f-h** and **<u>Supplementary</u> figure 7c-d)** indicating that alterations induced by acute inflammatory challenges were transient.

**Figure 4:**
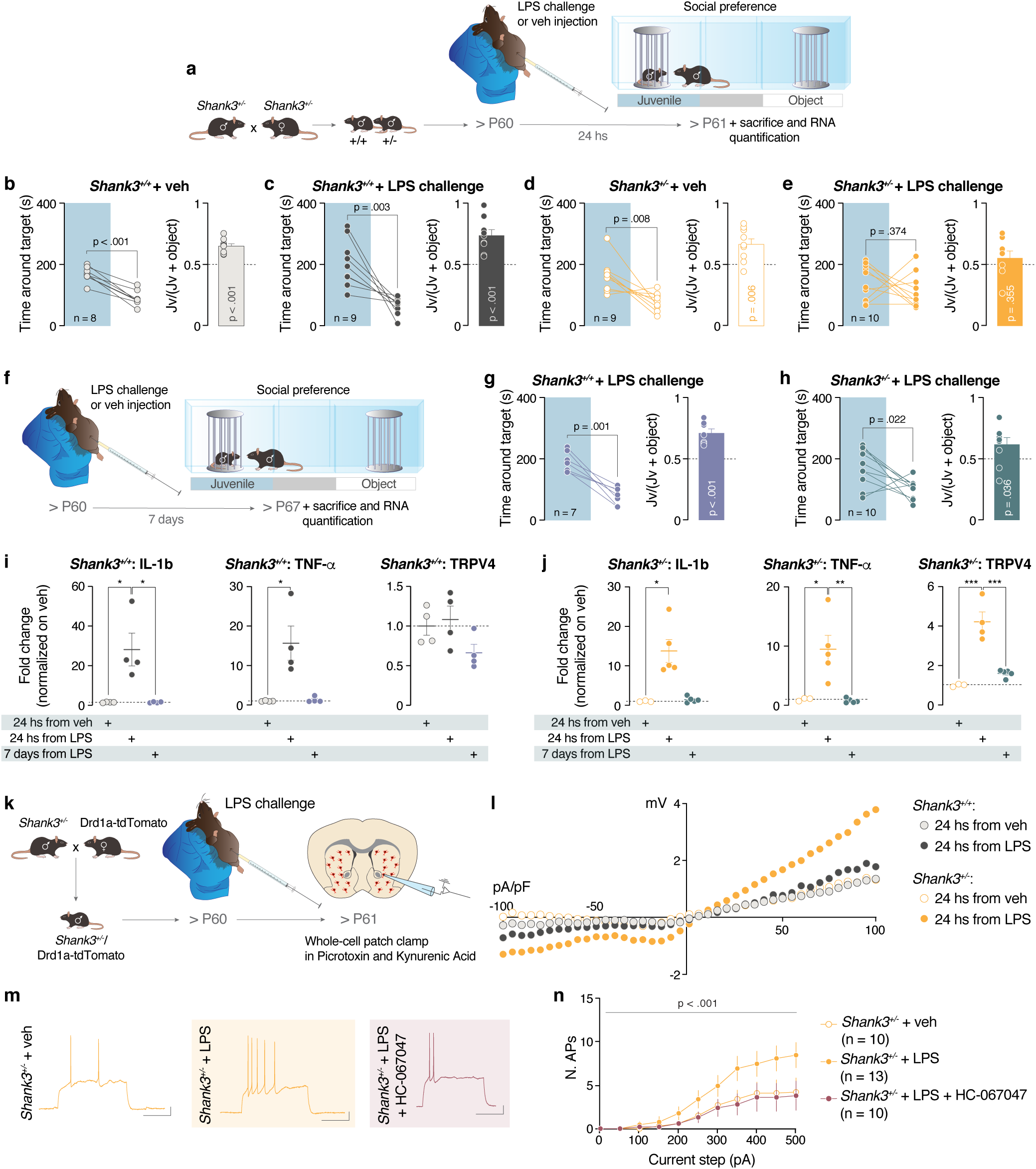
LPS challenge in *Shank3^+/-^* unmasks social deficits. **(a)** Experimental design. *Shank3^+/+^* and *Shank3^+/-^* were intraperitoneally injected with LPS or vehicle and 24 hrs later they were subjected to 3-chamber task. **(b, c, d, e)** Left: Time spent around the target (Paired-samples t-tests for object- vs. social: (b); *t* _(7)_ = 7.686, *p* < 0.001, (c); *t* _(8)_ = 4.199, *p* = 0.003, (d); *t* _(8)_ = 3.462, *p* = 0.009, (e); *t*_(9)_ = 0.935, *p* = 0.374). Right: juvenile preference index (one-sample t-tests against chance level = 0.5: (b); *t* _(7)_ = 7.2, *p* < 0.001, (c); *t* _(8)_ = 5.262, *p* < 0.001, (d); *t* _(8)_ = 3.734, *p* = 0.006, (e); *t* _(9)_ = 0.9747, *p* = 0.355). **(f)** Experimental design. *Shank3^+/+^* and *Shank3^+/-^* were intraperitoneally injected with LPS and 7 days later were subjected to 3-chamber task. **(g, h)** Left: Time spent around the target (Paired-samples t-tests for object- vs. social: (g); *t* _(6)_ = 5.979, *p* = 0.001, (h); *t* _(9)_ = 2.759, *p* = 0.022). Right: juvenile preference index (one-sample t-tests against chance level = 0.5: (g); *t* _(6)_ = 6.054, *p* < 0.001, (h); *t* _(9)_ = 2.463, *p* = 0.036). **(i)** mRNA expression analysis of *IL-1β*,*TNF-α* and *Trpv4* genes after LPS challenge in *Shank3^+/+^* (IL-1 β one way ANOVA followed by Sidak’s multiple comparisons test, *F_(2,9)_=10.33, p=0.005*; TNF-α Kruskal-Wallis statistic 7.538, *p=0.012; Trpv4* one way ANOVA followed by Sidak’s multiple comparisons test, *F_(2,9)_=2.768, p = 0.116* ). **(j)** mRNA expression analysis of *IL-1β*,*TNF-α* and *Trpv4* genes after LPS challenge in *Shank3^+/-^* (IL-1β Kruskal-Wallis statistic 9.002, *p=0.002*; TNF-α one way ANOVA followed by Sidak’s multiple comparisons test, *F_(2.10)_=10.27, p=0.004; Trpv4* one way ANOVA followed by Sidak’s multiple comparisons test: *F _(2,9)_ = 31.26, p < 0.001*). **(k)** Experimental design. *Shank3^+/-^* were crossed with Drd1a-tdTomato mice labelling specifically D1R-MSNs in *Shank3^+/-^* background. *Ex-vivo* patch clamp recordings were made 24 hrs after the LPS injection. **(l)** Whole-cell recording of Trpv4 current after LPS challenge in *Shank3^+/+-^* and *Shank3^+/-^* mice (Repeated measures ANOVA, voltage steps main effect *F _(1.974,43.29)_ =* 16.15, *p=0.001*, genotype by voltage steps interaction *F _(120 , 880)_ =* 1.451, *p =* 0.002; n = 5 cells, 2 mice (*Shank3^+/+^*), n=5 cells, 2 mice (*Shank3^+/+^*+LPS), n = 7 cells, 2 mice (*Shank3^-/+^*) n = 9 cells, 2 mice (*Shank3^-/+^*+LPS)) **(m)** Example traces from 300 pA depolarizing current injection in D1R+ MSNs of *Shank3^+/-^* mice after vehicle IP injection and treated with vehicle (left), D1R+ MSNs of *Shank3^+/-^* mice after LPS challenge and treated with vehicle (middle), and in D1R+ MSNs of *Shank3^+/-^* mice after LPS challenge and treated with HC-067047. **(n)** Number of action potentials (nAPs) across increasing depolarizing current steps (0-500 pA) for D1R-tom+:: *Shank3^+/-^* MSNs after LPS challenge (Repeated measures ANOVA, LPS challenge main effect *F _(2.30)_ =* 3.034, *p* = 0.063, current steps main effect *F _(10 , 300)_ =* 28.08, *p <* 0.001, LPS challenge by current steps interaction *F _(20 , 300)_ =* 2.042, *p =* 0.006, n = 10 cells, 4 mice (D1R-tom+:: *Shank3^+/-^* + veh), n = 13 cells, 3 mice (D1R-tom+:: *Shank3^+/-^* + LPS), n = 10 cells, 3 mice (D1R-tom+:: *Shank3^+/-^* + LPS + HC067047)). Error bars report SEM.

We next asked whether striatal *Trpv4* expression was altered in Shank3^+/-^ mice after LPS. Remarkably, whereas LPS injections increased the inflammatory markers IL-1β and TNF-α expression 24 hours after LPS injection in both *Shank3^+/+^* and *Shank3^+/-^* mice, the observed increase in *Trpv4* was seen only in *Shank3^+/-^* mice and not detectable 7 days after LPS challenge (**Fig. 4i-j**).

These data supported the hypothesis that inflammatory challenges may unmask behavioural phenotypes in *Shank3^+/-^* mice via an upregulation of *Trpv4* in the striatum.

### Hyperexcitability seen in D1R-MSNs of *Shank3^+/-^* mice after an acute LPS challenged is rescued by Trpv4 antagonist *ex-vivo*

To further investigate our hypothesis, we crossed *Shank3^+/-^* with Drd1a-tdTomato mice and we performed *ex-vivo* patch-clamp recordings from D1R-MSNs 24 hrs after LPS injection (**Fig 4k**). In order to probe the functional consequences of *Trpv4* upregulation, we first assessed Trpv4-mediated whole-cell currents from D1R-MSNs and observed an increase only in LPS-challenged *Shank3^+/-^* mice (**Fig 4l**). Importantly, we found that LPS challenge caused a D1R-MSNs hyperexcitability in *Shank3^+/-^* mice similarly to the NAc-sh*Shank3* model (**Fig 4m-n**). Finally, bath application of Trpv4 antagonist, HC-067047, normalized neuronal excitability, strengthening the causal links between D1R-MSN hyperexcitability and *Trpv4* upregulation (**Fig 4m-n**).

### Intra-NAc Trpv4 antagonist restores sociability in *Shank3^+/-^* LPS-challenged mice

Although these results further corroborated our hypothesis, we still questioned whether the NAc plays a role in the behavioural alterations observed in *Shank3^+/-^* mice after LPS injection.

To answer this question, *Shank3^+/-^* mice were bilaterally cannulated above the NAc for local pharmacology experiments. After one week of recovery, mice were treated with either HC-067047 or vehicle intra-NAc infusions 1 hour after the LPS challenge. The day after, mice were again infused locally in the NAc with HC-067047 or vehicle 30 minutes before the three-chamber task (**Fig. 5a**). While *Shank3^+/-^*::LPS mice infused with vehicle showed sociability deficits (**Fig. 5b** and **Supplementary figure 8a**), *Shank3^+/-^*::LPS mice infused with HC-067047 spent more time around the enclosure containing the juvenile mouse (**Fig. 5c**), albeit without a significant difference in the time spent in chambers (**Supplementary figure 8a**). Locomotor activity was not affected by local HC-067047 treatment (**Supplementary figure 8b**). These results indicate that the inhibition of *Trpv4* in the NAc after immune system activation is sufficient to ameliorate sociability deficits, suggesting a link between *Trpv4* modulation and social behaviour.

**Figure 5:**
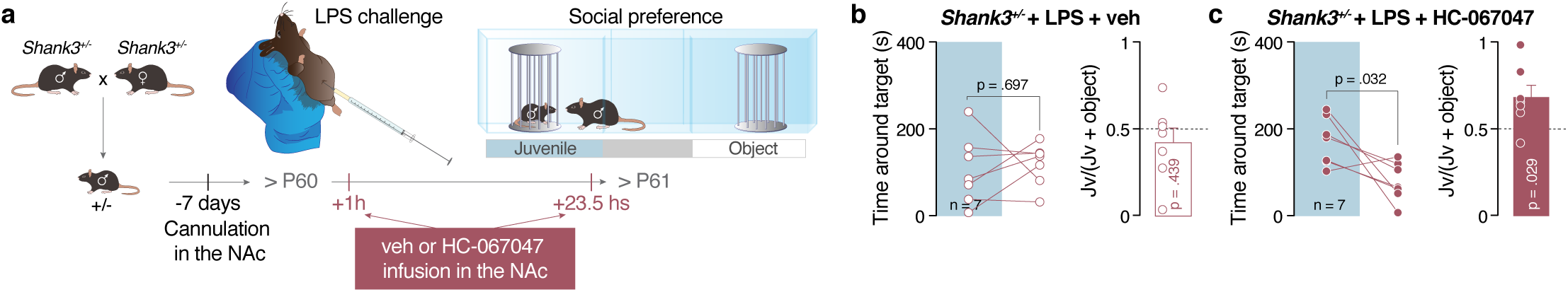
Trpv4 antagonist infused in the NAc improves social deficits in *Shank3^+/-^* mice challenged with LPS. **(a)** Experimental design. Adult *Shank3^+/-^* mice were intraperitoneally injected with LPS and 24 hours later were subjected to the behavioural task. 30 minutes before the test, mice were infused (in the NAc) either with Trpv4 antagonist (HC-067047) or vehicle. **(b, c)** Left: Time spent around the target for *Shank3^+/^* mice after LPS challenge and vehicle or HC-067047 infusion in the NAc (Paired-samples t-tests for object- vs. social: (b); *t* _(6)_ = 0.408, *p* = 0.687, (c); *t* _(6)_ = 2.787, *p* = 0.032). Right: juvenile preference index (one-sample t-tests against chance level = 0.5: (b); *t* _(6)_ = 0.629, *p* = 0.4388, (c); *t* _(6)_ = 2.852, *p* = 0.029). Error bars report SEM.

Collectively, our data highlight the NAc *Trpv4* alterations as a potentially common and unifying molecular underlying factor in sociability and aberrant intrinsic neuronal properties in *Shank3* mouse models for autism.

## DISCUSSION

Mutations in the *SHANK3* gene have been recognized as a genetic risk factor for ASD. Remarkably, high heterogeneity of neuronal pathophysiology and behavioral phenotypes have been reported in *Shank3* mouse models. Nevertheless, whether environmental factors contribute to the phenotypic heterogeneity of *Shank3* mouse model is still largely unknown. Here we first found that early loss of *Shank3* in the Nac reduces sociability via direct pathway hyperexcitability. These changes were accompanied by an unbalance of inflammatory mediators and by the overexpression of transient receptor potential vanilloid 4 (*Trpv4*). Interestingly, lipopolysaccharide-induced neuroinflammation revealed similar molecular, circuit and behavioural alterations in genetically vulnerable *Shank3*^+/-^ mice. Acute Trpv4 inhibition in the NAc restored excitability and sociability deficits. Our data not only suggest that activation of the immune system may unmask autism-related behavioural phenotypes in genetically vulnerable mice but also ascribe Trpv4 as a potential therapeutic target for sociability defects in Autism.

The mesolimbic system represents an interesting hub for ASD pathophysiology. Indeed, human studies reported that social stimuli activate the NAc^35–39^ and that this activation is disrupted in ASD patients^11, 40^. In support of the clinical studies, alterations in the mesolimbic system induce reward-related behavioural alterations in rodents^8, 13, 41^. However, the neuronal mechanisms underlying NAc-related sociability deficits remained largely unknown. It has been previously shown that the lack of *Shank3* induces differential alterations of intrinsic and synaptic properties of dorsolateral striatum D1R- and D2R-MSNs and that deficits in the indirect pathway contribute to repetitive behaviour^3, 6^. In our study, we found that the downregulation of *Shank3* in the ventral striatum alters sociability via hyperexcitability of D1R-MSNs, which is linked to gene expression alterations. While we cannot exclude that changes in D2R-MSNs also contribute to the phenotype, it is important to note that changes in excitability in the indirect pathway neurons were only observed in absence of synaptic blockers. Furthermore, while decreasing the activity of D1R-MSNs in sh*Shank3* mice was able to rescue the behavioural phenotype, decreasing the activity of the direct pathway neurons in control mice alters sociability (**Fig. 2h**). These data not only causally link the activity of the direct pathway ventral striatum to sociability but suggest that the activity of D1R-MSN has to be tightly tuned in order to guarantee optimal expression of social behaviour. These findings are in line with previous evidence supporting the importance of NAc D1R-MSNs activity in modulating social behavior^25^.

Although ASD is known as a synaptic pathology^46, 47^, recent evidence demonstrated a fundamental role of ion channels deficits in the pathophysiology of ASDs. Indeed, the loss of scaffolding between Shank3 and HCN impairs Ih currents and neuronal excitability^28^. Here, we highlighted a novel link between Trpv4 alterations and Shank3 insufficiency. We found that accumbal *Shank3* insufficiency upregulates *Trpv4*, a non-selective cation channel constitutively active at physiological temperatures^42^, which allows Ca^2+^ influx, stimulates Ca^2+^-induced Ca^2+^- release (CICR) signaling^43–45^ and ultimately tunes neuronal excitability^42^. Interestingly, we have not observed an upregulation of *Trpv4* in P90-injected sh*Shank3* mice. To further prove the causal link between the gene and behaviour, we observed an increase in Trpv4 expression in *Shank3^+/-^* mice 24 hrs after LPS, time point at which we also could observe behavioural deficits. Furthermore, sociability of sh*Shank3* and LPS-*Shank3^+/-^* mice improves by a region-specific Trpv4 inhibition. Although future experiments will have to determine the precise mechanisms of how a scaffold protein could affect the transcription of a set of genes, our study supports the idea that *Shank3* downregulation affects both intrinsic excitability and synaptic properties, which may ultimately account for the symptom heterogeneity of PMS patients.

The heterogeneity of ASD symptoms most likely results from the involvement of a multitude of genetic factors and a complex interaction between those genes and environmental challenges^48–53^. For example, increasing evidence suggests a role for inflammation in ASD pathogenesis^54–56^. Indeed, individuals with ASD often have heightened levels of pro-inflammatory cytokines^57, 58^, and post mortem brain samples revealed an upregulation of genes related to the immune response^59^. A recent hypothesis posits that the activation of the immune system during critical periods of brain development may cause neuronal dysfunctions^60, 61^ and lead to behavioural deficits^62^. To better understand how immune responses to infectious agents might affect behavior in preclinical models, we used LPS, a bacterial endotoxin, that stimulates an innate response to bacterial infection leading to a variety of behavioural changes^63–68^. Interestingly, animals exposed to inflammatory stimuli show impaired motivation, decreased exploratory behaviour^69^, and social withdrawal^70, 71^. Here, we show that one LPS injection during adulthood reveals transient sociability deficits in adult *Shank3^+/-^* mice. Our data suggest that immune system activation may expose an underlying genetic vulnerability in *Shank3^+/-^* mice, leading to social behavior deficits.

Exploring the contribution of striatal dysfunctions to ASD pathophysiology allowed us to uncover alterations in specific neuronal populations and to find a novel potential therapeutic target. Specifically, using a circuit-specific knock-down strategy, we identified *Trpv4* upregulation as the link between changes in excitability, inflammatory response and behavioural deficits. Trpv4 is widely expressed in the brain where it is activated by changes in both osmotic pressure and heat^72–74^. Research into the involvement of Trpv4 in neuropathies and neurodegenerative diseases has attracted an increasing interest^75, 76^. Indeed, whole-genome sequencing of quartet families with ASD has revealed frameshift mutations of *Trpv4*^77^, suggesting a possible involvement in the pathogenesis of autism. Furthermore, hyperactivity of these channels occurs in several pathological conditions^45, 76, 78, 79^. Interestingly, Trpv4 activation may induce inflammation by increasing pro-inflammatory cytokines^45, 80^ and Trpv4 inhibitors have been used to counteract oedema and inflammation^31^. Although it is well established that inflammatory cytokines may impact both synaptic transmission and neuronal excitability^81, 82^, the direct link between Trpv4, neuronal function and behaviour was still relatively unknown. Here, using a circuit approach, we firstly identify an upregulation of *Trpv4* after *Shank3* downregulation. Consequently, based on these results, we found that inflammatory challenge in *Shank3^+/-^* mice increased the expression of *Trpv4* and induced D1R-MSNs hyperexcitability. By rescuing sociability deficits in these mice, we provide a novel link between immunoresponse, genetic background, and neuronal activity in the context of ASD. Finally, our data point at Trpv4 channel as a novel potential candidate for the treatment of ASD symptoms.

Overall, our data highlight that viral-mediated and region-specific ablation of *Shank3*, is a suitable model to obtain mechanistic insights regarding regions and cell types that could be implicated in autism-relevant symptoms and furthermore, to validate hypotheses and potential novel therapeutic interventions.

## Authors Contributions

ST, SM, AC and CB conceived and designed the experiments. ST, AC and GC performed and analyzed the behavioural experiments. ST, SM and SB performed and analyzed the electrophysiological experiments. EM and DJ analyzed the results obtained from the bulk RNA sequencing. YJ generated the mutated Shank3 mouse line. ST, SM, AC and CB wrote the manuscript and AC prepared the figures.

## Acknowledgments

CB is supported by the Swiss National Science Foundation, Pierre Mercier Foundation, ERC consolidator grant and NCCR Synapsy. We thank Lorena Jourdain for technical support.

## Figure legends

**Supplementary Figure 1:**
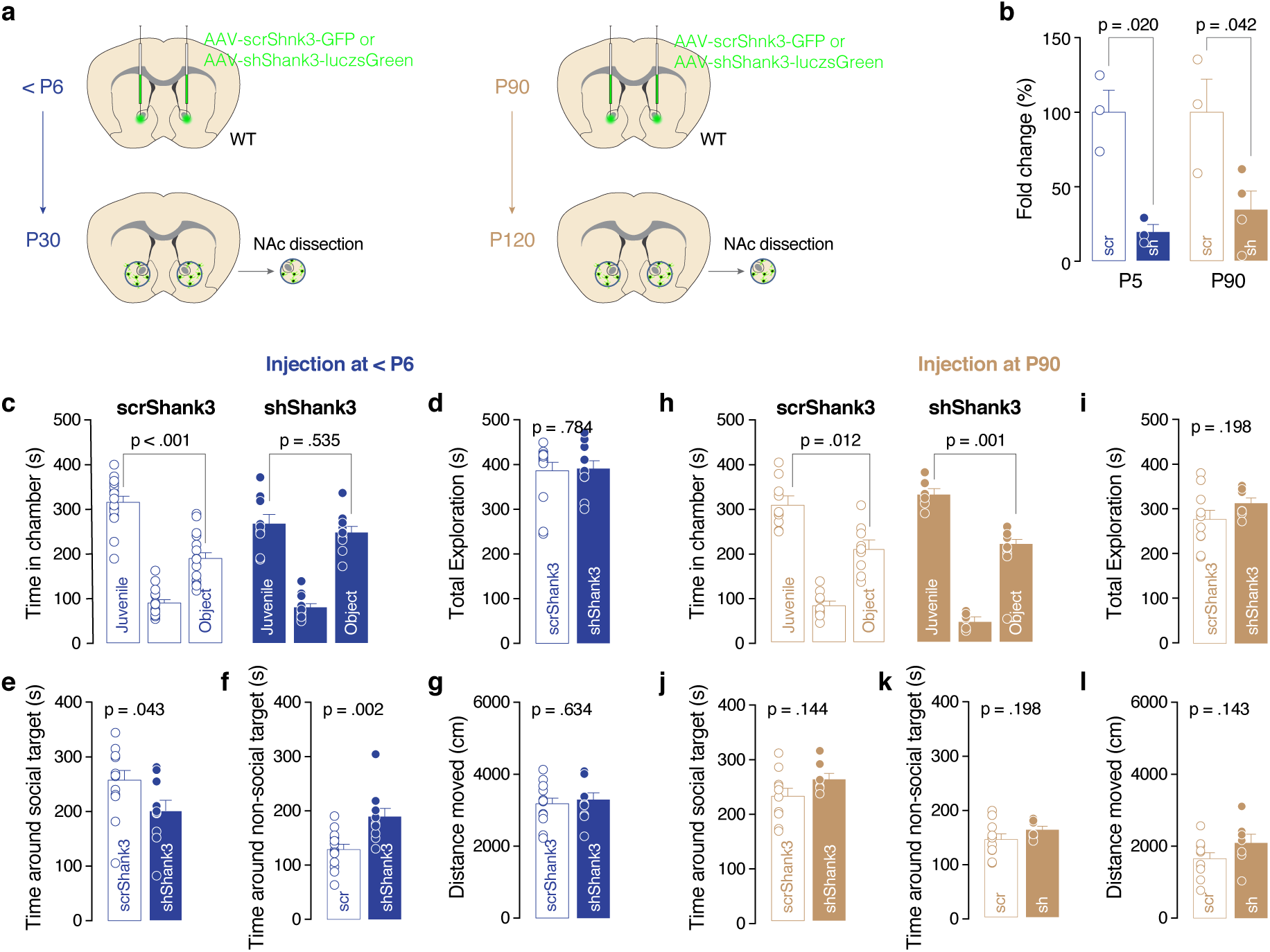
*Shank3* downregulation during development increases the interaction with the non-social target. **(a)** Schema of injection sites in the NAc with AAV-scrShank3-GFP or AAV-shShank3-luczsGreen in ≤P6 or P90 mice. Subsequently, the NAc was dissected and mRNA was extracted. **(b)** Real-time PCR analysis of NAc dissected from P6- or P90-injected mice confirm the downregulation of *Shank3* in sh infected mice (two-way ANOVA followed by Bonferroni’s multiple comparisons test: Virus main effect *F_(1, 8)_ = 18.80, p=0.003*). **(c, h)** Time spent in compartments for mice injected ≤ P6 ((c) Paired-samples t-tests for object- vs. social-containing chambers: scrShank3; *t* _(12)_ = - 5.047, *p* < 0.001), shShank3; *t* _(9)_ = - 0.645, *p* =0.535, (h) Paired-samples t-tests for object- vs. social-containing chambers: scrShank3; *t* _(9)_ = 3.144, *p* = 0.012, shShank3; *t* _(6)_ = 5.686, *p* = 0.001). **(d)** Total exploration time around the enclosures for mice injected neonatally (Mann-Whitney test, *p* = 0.784). **(e)** Time spent around the enclosure containing the social stimulus (*t* _(21)_ = 2.152, *p* = 0.043). **(f)** Time spent around the non-social target (*t* _(21)_ = -3.499, *p* = 0.002). **(g)** Distance moved in the apparatus (*t*_(21)_ = -0.483, *p* = 0.634). **(i)** Total exploration time around the enclosures for mice injected during adulthood (*t* _(15)_ = 1.347, *p* = 0.198). **(j)** Time spent around the enclosure containing the social stimulus (*t* _(15)_ = -1.541, *p* = 0.144). **(k)** Time spent around the non-social target (*t* _(15)_ = 1.347, *p* = 0.198). **(l)** Distance moved in the apparatus (*t* _(15)_ = -1.544, *p* = 0.143). Error bars report SEM.

**Supplementary Figure 2:**
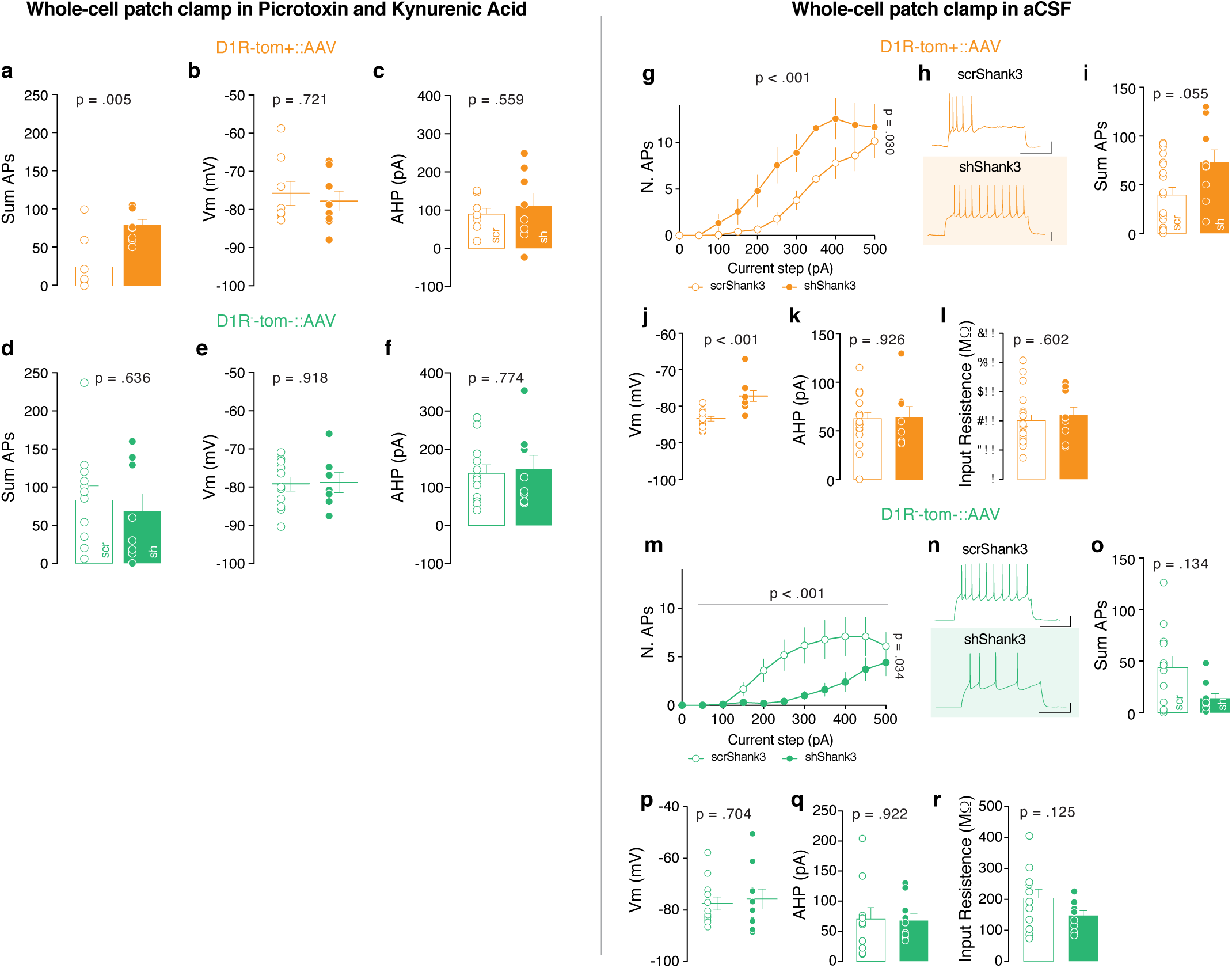
*Shank3* NAc downregulation alters D1R MSNs excitability. **(a, d, i, o)** Total number of APs across all steps ((a) Mann Whitney test, *p =* 0.005. (d) unpaired t-test, *t _(18)_ =* 0.482 *p =* 0.636. (i) Mann Whitney test, *p =* 0.055. (o) Mann Whitney test, *p =* 0.134.). **(b, e, j, p)** Resting membrane potential of recorded cells ((b) Mann Whitney test, *p =* 0.721. (e) unpaired t-test, *t_(17)_ =* 0.105 *p =* 0.918. (j) Mann Whitney test, *p <* 0.001. (p) unpaired t-test, *t* _(20)_ = 0.385 *p =* 0.704). **(c, f, k, q)** After-hyperpolarization current (AHP) of recorded cells ((c) unpaired t-test, *t _(14)_ =* 0.597, *p=*0.559. (f) unpaired t-test, *t _(18)_ =* 0.291 *p =* 0.774. (k) unpaired t-test, *t _(24)_* = 0.094*, p =* 0.926. (q) unpaired t-test, *t* _(18)_ = 0.099 *p =* 0.922). **(g)** Number of action potentials (nAPs) across increasing depolarizing current steps (0-500 pA) for D1R-tom+::scrShank3 and shShank3 MSNs (Repeated measure (RM) two-way ANOVA, main effect of virus *F _(1, 27)_* = 5.285 *p =* 0.030, main effect of current steps *F _(10, 270)_ =* 32.46 *p <* 0.001, virus by current steps interaction *F _(10 , 270)_ =* 1.957 *p =* 0.038, n = 9 cells, 3 mice (shShank3), n = 20 cells, 5 mice (scrShank3)). **(h)** Example traces from 300 pA depolarizing current injection in D1R-tom+ MSNs infected with scrShank3 (upper part) or with shShank3 (lower part). **(l, r)** Input resistance of recorded cells ((l) *t _(27)_ =* 0.528, *p =* 0.602. (r) *t* _(19)_ = 1.607, *p* = 0.125). **(m)** Number of action potential (nAPs) across increasing depolarizing current steps (0-500 pA) for D1R-tom-::scrShank3 and shShank3 MSNs (Repeated measures ANOVA, main effect of virus *F _(1, 20)_ =* 5.207, *p =* 0.034, main effect of current steps *F _(10, 200)_ =* 11.77, *p <* 0.001, virus by current steps interaction *F _(10, 200)_ =* 2.958 *p* = 0.002, n = 10 cells, 3 mice (shShank3), n=12 cells, 3 mice (scrShank3)). **(n)** Example traces from 300 pA depolarizing current injection in D1R-tom-MSNs infected with scrShank3 (upper part) or with shShank3 (lower part). Error bars report SEM.

**Supplementary Figure 3:**
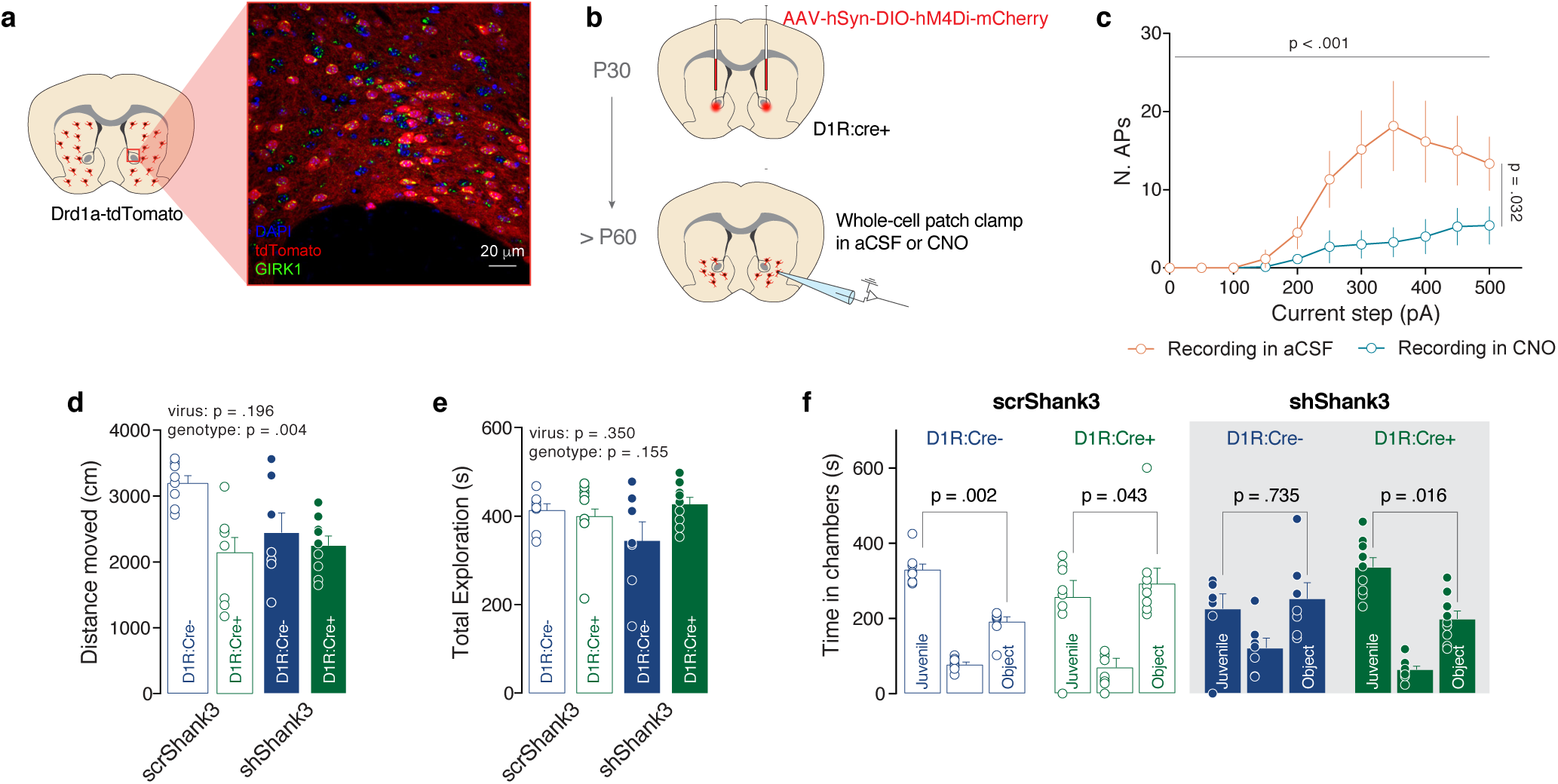
Dampening D1R-MSNs activity improves social deficits in NAc-shShank3 mice. **(a)** Representative image of GIRK1 expression (green) in the NAc of Drd1a-dTomato (red) mice. **(b)** Experimental design. D1R-Cre positive (D1R:Cre^+^) mice were injected in the NAc with AAV-hSyn-DIO-hM4Di-mCherry (DREADD) and after P60 whole-cell patch clamp recordings were performed. NAc slices were either pre-incubated with CNO and recorded in presence of CNO, or were incubated and recorded in aCSF only. **(c)** Number of action potentials (nAPs) across increasing depolarizing current steps (0-500 pA) in presence or absence of CNO. The number of APs was significantly decreased by the bath application of CNO (Repeated measures ANOVA, main effect of drug *F _(1, 11)_ =* 6.060 *p* = 0.032, main effect of current steps *F _(10, 110)_ =* 12.11 *p <* 0.001, drug by current steps interaction *F _(10, 110)_* = 4.342 *p* < 0.001, n = 6 cells (aCSF) 7 cells (CNO); n = 2 mice). **(d)** Distance moved for D1R:Cre+/- mice injected with DREADD and scrShank3 or DREADD and shShank3 (Two-way ANOVA: main effect of virus *F* _(1, 28)_ = 1.756, *p* = 0.196, main effect of genotype *F* _(1, 28)_ = 10.039, *p* = 0.004, virus by genotype interaction *F* _(1, 28)_ = 4.959, *p* = 0.034). **(e)** Total exploration time for D1R:Cre^+^ or D1R:Cre^-^ mice injected with DREADD and scrShank3 or DREADD and shShank3 (Two-way ANOVA: main effect of virus *F* _(1, 28)_ = 0.902, *p* = 0.350, main effect of genotype *F* _(1, 28)_ = 2.135, *p* = 0.155, virus by genotype interaction *F* _(1, 28)_ = 2.604, *p* = 0.118). **(f)** Time spent in compartments of the three-chamber social interaction task for D1R:Cre^+^ or D1R:Cre^-^ mice injected with DREADD and scrShank3 or DREADD and shShank3 (D1R:Cre^-^::scrShank3: *t* _(7)_ = 4.916, *p* = 0.002, D1R:Cre^+^::scrShank3: *t* _(7)_ = 0.043, *p* = 0.967, D1R:Cre^-^::shShank3: *t* _(6)_ = -0.355, *p* = 0.735, D1R:Cre^+^::shShank3: *t* _(8)_ = 3.031, *p* = 0.016). Error bars report SEM.

**Supplementary Figure 4:**
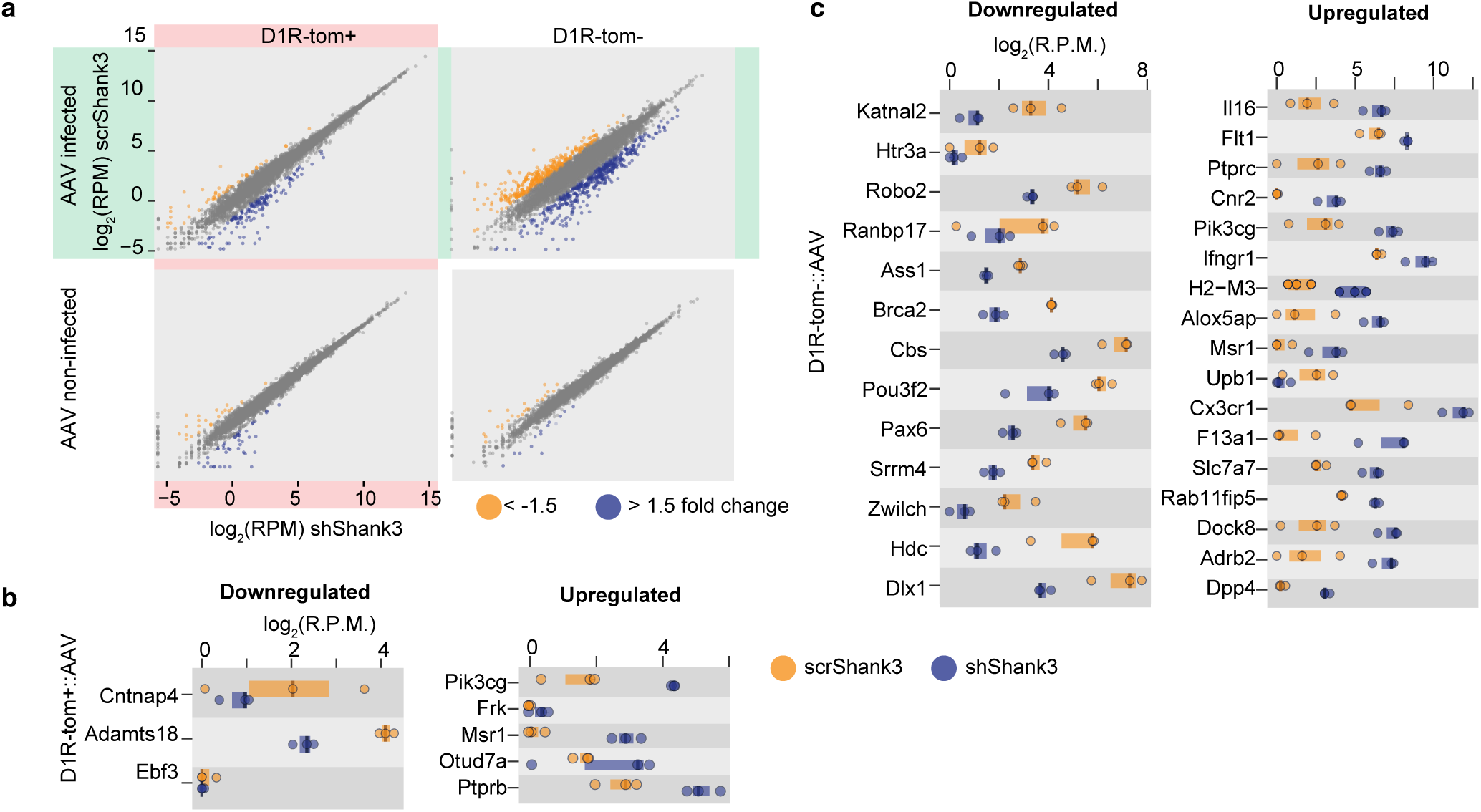
D1R-MSNs shShank3 downregulated genes association with SFARI genes. **(a)** Differential expression analysis of AAV-scrShank3 vs AAV-ShShank3 shows small indirect transcriptional effect in non-infected samples, while infected samples display the stronger transcriptomic alterations. In both D1R+ **(b)** and D1R- **(c)** SFARI associated genes are altered, supporting the link between Shank3 downregulation with an autism-related phenotype.

**Supplementary Figure 5:**
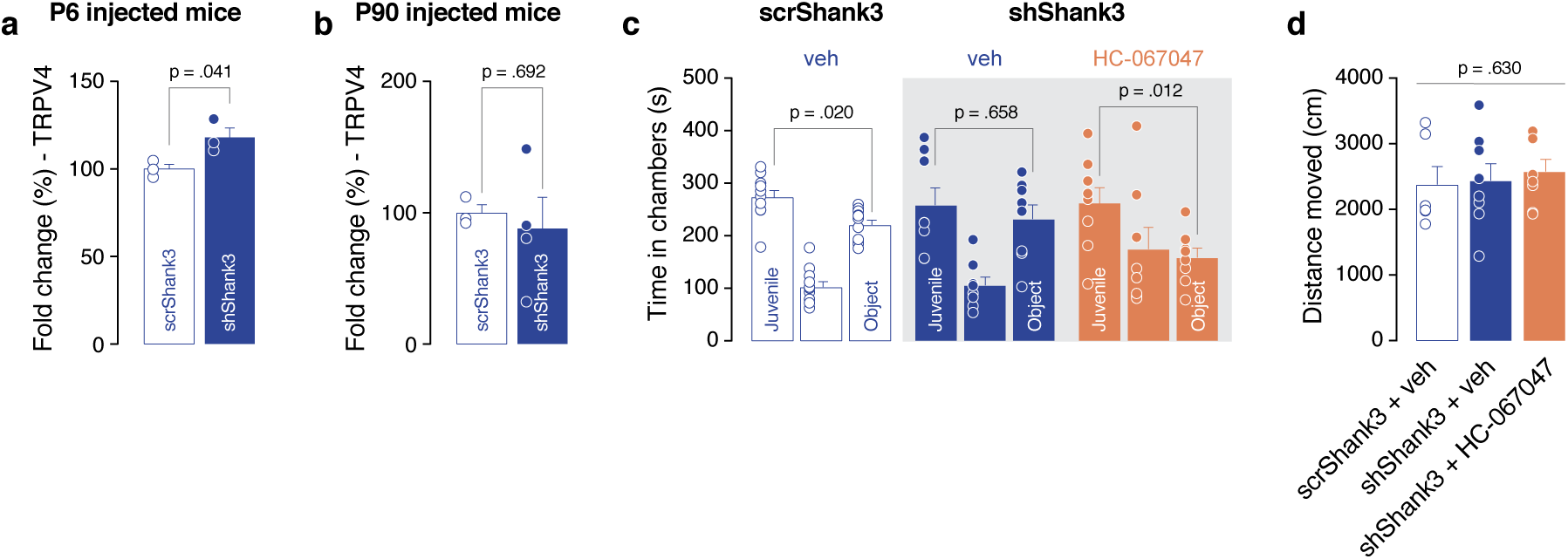
Supplementary data on behavioral experiments showed in Figure 4. **(a-b)** Real-time PCR analysis of NAc dissected from P6- or P90-injected mice confirm the upregulation of *Trpv4* in P6 sh-infected mice (P6, unpaired t-test *t* _(4)_ = 2.980, *p* = 0.041, P90 *t* _(5)_ = 0.4203, *p* = 0.69). **(c)** Time spent in compartments for mice infected with scrShank3 or shShank3 and infused with vehicle or HC-067047 (Paired-samples t-tests for object- vs. social-containing chambers scrShank3 + veh: *t* _(10)_ = 2.772, *p* = 0.020, shShank3 + veh: *t* _(7)_ = 0.462, *p* = 0.658, shShank3 + HC-067047: *t* _(7)_ = 3.339, *p* = 0.012). **(d)** Distance moved during social preference test (one way ANOVA followed by Bonferroni’s multiple comparisons test: *F*_(3,24)_=0.586, *p* = 0.630). Error bars report SEM.

**Supplementary Figure 6:**
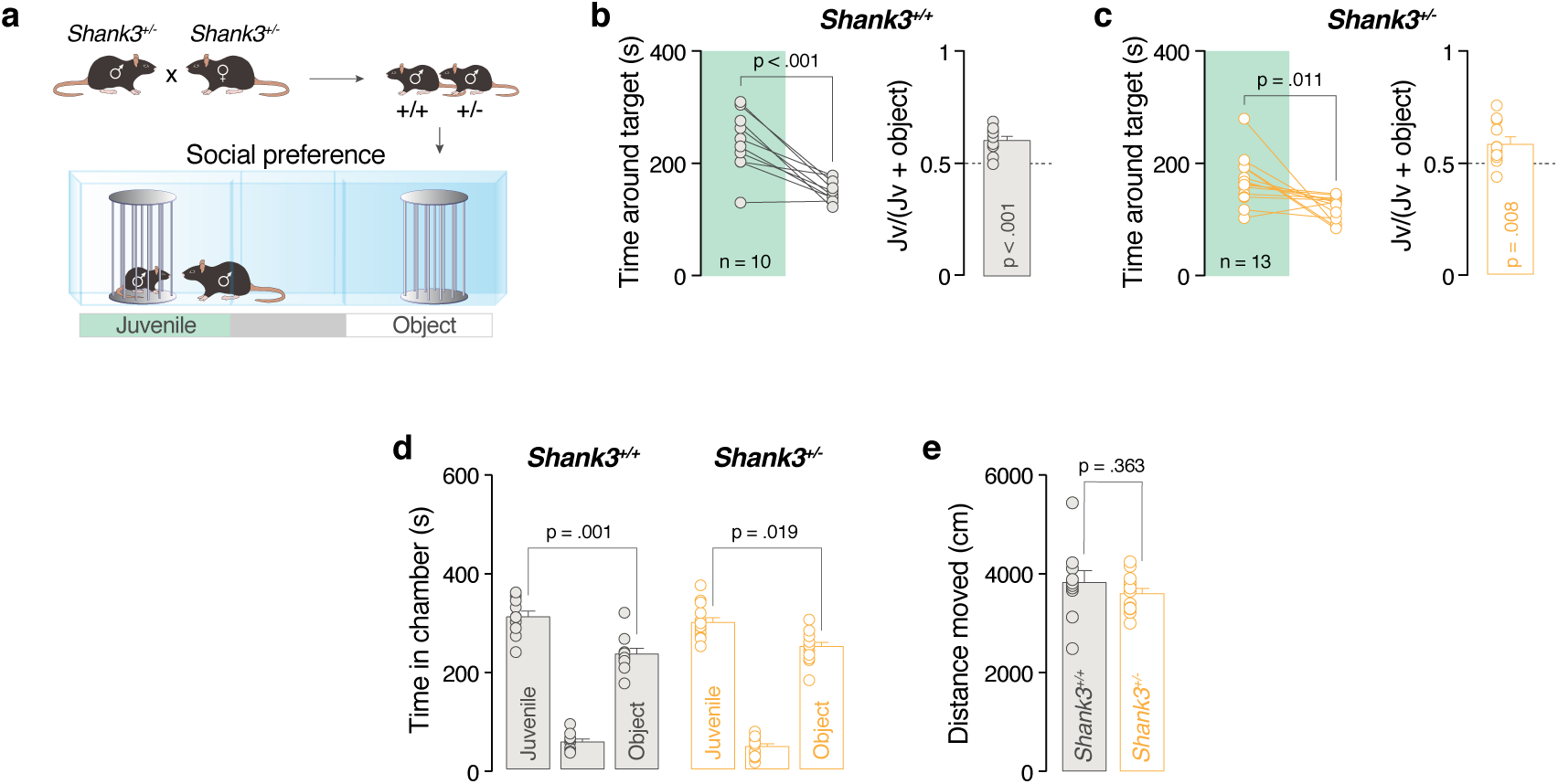
*Shank3^+/-^* does not show social deficits. **(a)** Behavioural task paradigm. **(b, c)** Left: Time spent around the target during social preference test for *Shank3^+/+^* and *Shank3^+/-^* mice (paired-samples t-tests for object- vs. social: *Shank3^+/+^*: *t* _(9)_ = 5.167, *p* < 0.001; *Shank3^+/-^*: *t* _(12)_ = 3.026, *p* = 0.011). Right: juvenile preference index (one-sample t-tests against chance level = 0.5: *Shank3^+/+^*: *t* _(9)_ = 5.617, *p* < 0.001; *Shank3^+/-^*: *t* _(12)_ = 3.146, *p* = 0.008). **(d)** Time spent in the juvenile, object or in center chamber during social preference test for *Shank3^+/+^*, *Shank3^+/-^* and *Shank3^-/-^* mice (Paired-samples t-tests for object- vs. social-containing chambers: *Shank3^+/+^*: *t* _(9)_ = 3.269, *p* = 0.009; *Shank3^+/-^*: *t* _(12)_ = 2.705, *p* = 0.019). **(e)** Distance moved during social preference test (Unpaired-samples t-tests: *t* _(21)_ = 0.9307, *p* = 0.363).

**Supplementary Figure 7:**
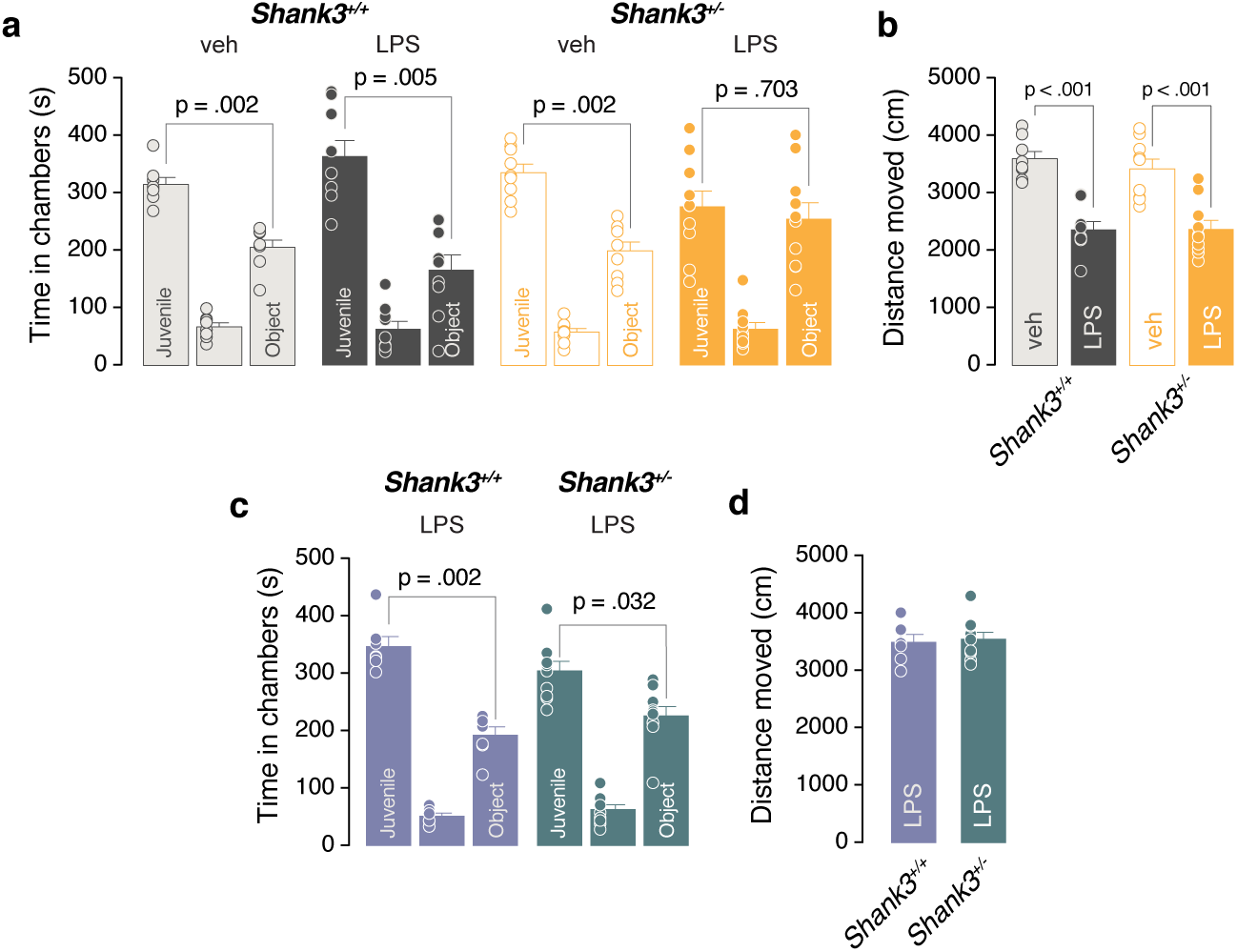
Supplementary data on behavioral experiments showed in Figure 5. LPS challenge induces social deficits in *Shank3^+/-^* mice after 24 hours: **(a)** Time spent in the juvenile, object or in the center chamber during social preference test for *Shank3^+/+^* and *Shank3^+/-^* previously injected with vehicle or LPS (Paired-samples t-tests for object- vs. social-containing chambers: *Shank3^+/+^* + veh: *t* _(7)_ = 4.838, *p* = 0.002, *Shank3^+/+^* + LPS: *t* _(8)_ = 3.87, *p* = 0.005, *Shank3^+/-^* + veh: *t* _(8)_ = 4.526, *p* = 0.002, *Shank3^+/-^* + LPS: *t* _(9)_ = 0.3939, *p* = 0.703). **(b)** Distance moved during social preference test (two-way ANOVA followed by Bonferroni’s multiple comparisons test: LPS treatment main effect *F_(1,32)_=58.03, p<0.001*). 7 days after LPS challenge the sociability and the distance moved are not impaired anymore: **(c)** Time spent in the juvenile, object or in the center chamber during social preference test for *Shank3^+/+^* and *Shank3^+/-^* previously injected LPS (Paired-samples t-tests for object- vs. social-containing chambers: *Shank3^+/+^* + LPS: *t* _(7)_ = 5.083, *p* = 0.002, *Shank3^+/-^* + LPS: *t* _(10)_ = 2.536, *p* = 0.032. **(d)** Distance moved during social preference test (unpaired t-test t*_(15)_*=0.3116, p=0.76). Error bars report SEM.

**Supplementary Figure 8:**
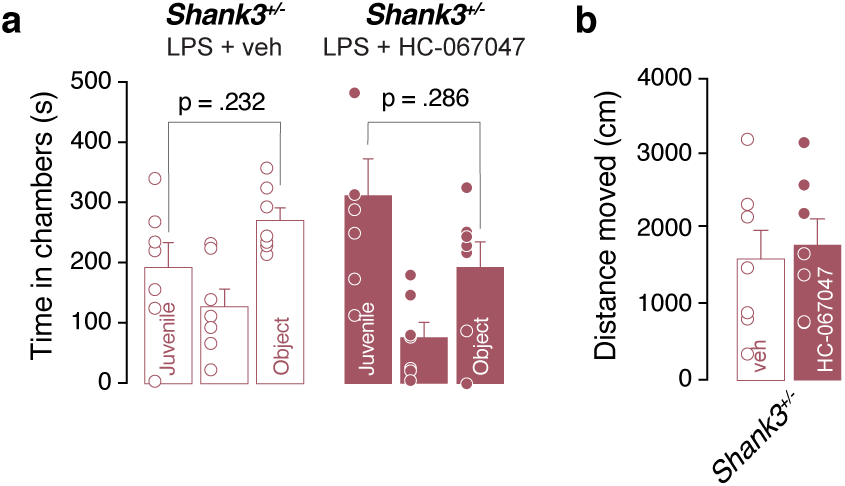
Supplementary data on behavioral experiments showed in Figure 6. **(a)** Time spent in compartments for *Shank3^+/-^* mice after LPS challenge and with vehicle or HC-067047 infusion in the NAc (Paired-samples t-tests for object- vs. social-containing chambers: *Shank3^+/-^* + veh: *t* _(6)_ = 1.3229, *p* = 0.232, *Shank3^+/-^* + HC-067047: *t* _(6)_ = 1.17, *p* = 0.286). **(b)** Distance moved during social preference test (unpaired t-test: *t* _(12)_ = 0.3708, *p* = 0.717). Error bars report SEM.

## METHOD DETAILS

### Viruses and stereotactic injections

Viruses used in this study: (1) purified scr*Shank3* and sh*Shank3* (AAV1-GFP-U6-scrmbshRNA; titer: 5.9×10^13^ GC/mL and AAV5-ZacF-U6-luczsGreen-sh*Shank3*; titer: 7.4×10^13^ GC/mL, VectorBioLab); (2) AAV5/hsyn-DIO-hM4D(Gi)-mCherry (AV44961, titer: 5.5×10^12^ virus molecules/mL, UNC GTC vector core). Viral injections in the NAc were delivered in mice either at an early time-point (at P5 or P6; ≤P6) or later in life (>P30) (, depending on the experimental cohort. After anesthesia induction with a mixture of isoflurane/O2, C57Bl/6j wildtype pups or >P30 mice were placed on a stereotaxic frame (Angle One; Leica, Germany). For the pups, the coordinates used were AP: +3.5 mm, ML: ±0.8 mm, DV: -3.2 mm (measured from lambda), and for >P30 mice, the coordinates were AP: +1.2 mm, ML: ±1.0 mm, DV: -4.4/-4.0 mm (measured from bregma). To obtain bilateral NAc infection, 100 nl of viral solution was infused per injection in pups and 150 nl of viral solution was infused per injection in >P30 mice.

### Social preference test

A three-chambered social interaction assay was used, comprising a rectangular Plexiglas arena (60 × 40 × 22 cm) (Ugo Basile, Varese, Italy) divided into three chambers (each 20 × 40 × 22 (h) cm). The walls of the center chamber had doors that could be lifted to allow free access to all chambers. The social preference test was performed similarly as published by Moy *et al* ^83^. Briefly, each mouse was placed in the arena for a habituation period of 10 min, when it was allowed to freely explore the empty arena. At the end of the habituation, the test was performed: two enclosures with metal vertical bars were placed in the center of the two outer chambers. One enclosure was empty (serving as an inanimate object) whereas the other contained a social stimulus (unfamiliar juvenile mouse 25 ± 1 day old). The enclosures allowed visual, auditory, olfactory, and tactile contact between the experimental mice and the mice acting as social stimuli. The juvenile mice in the enclosures were habituated to the apparatus and the enclosures for a brief period of time on the 3 days preceding the experiment. The experimental mouse was allowed to freely explore the apparatus and the enclosures for 10 min. The position of the empty vs. juvenile-containing enclosures alternated and was counterbalanced for each trial to avoid any bias effects. Animals that their total exploration time for both the enclosures was less than 10 seconds were excluded from the analysis. In particular, one mouse in the D1:Cre^+^::scr*Shank3* group was excluded from the analysis according to this criterion. Every session was video-tracked and recorded using Ethovision XT (Noldus, Wageningen, the Netherlands), which provided an automated recording of the time around the enclosures (with virtual zones designed around them), the distance moved and the velocity. The time spent around each enclosure was assessed and then used to determine the preference score for the social target as compared to the empty enclosure (social/(social + empty)). The arena was cleaned with 1% acetic acid solution and dried between trials.

In the rescue experiment with the chemogenetic approach, 30 minutes before the habituation, scr- and sh*Shank3* injected mice were intraperitoneally injected with Clozapine N-oxide (CNO, Cat. No.: BML-NS105-0025, Lot No.: 07131709) dissolved in saline (5 mg/Kg).

In the LPS challenge experiments, *Shank3^+/+^* and *Shank3^+/-^* mice were intraperitoneally injected 24 hours before the test with LPS at a dose of 2mg/Kg in saline (NaCl 0.9%) (Lipopolysaccharides from Escherichia coli O26:B6, Sigma-Aldrich)

### Whole-cell patch clamp recordings

Coronal midbrain slices 250 μm thick containing the NAc were prepared following the experimental injection protocols described above. Brain were sliced in artificial cerebrospinal fluid (aCSF) containing 119 mM NaCl, 2.5 mM KCl, 1.3 mM MgCl_2_, 2.5 mM CaCl_2_, 1.0 mM NaH_2_PO_4_, 26.2 mM NaHCO_3_ and 11 mM glucose, bubbled with 95% O_2_ and 5% CO_2_. Slices were kept for 20-30 min at 35°C and then transferred at room temperature. Whole-cell voltage clamp or current clamp electrophysiological recordings were conducted at 32°–34° in aCSF (2–3 ml/min, submerged slices). Recording pipette contained the following internal solution: 140 mM K-Gluconate, 2 mM MgCl2, 5 mM KCl, 0.2 mM EGTA, 10 mM HEPES, 4 mM Na2ATP, 0.3 mM Na3GTP and 10 mM creatine-phosphate. The cells were recorded at the access resistance from 10–30 MΩ. Resting membrane potential (in mV) was read using the Multiclamp 700B Commander (Molecular Devices) while injecting no current (I = 0) immediately after breaking into a cell. Action potentials (AP) were elicited in current clamp configuration by injecting depolarizing current steps (50 pA, 500 ms) from 0 to 500 pA, in presence of Picrotoxin (100µM) and Kynurenic acid (3mM). For CNO validation and HC-067047 rescue, slices were incubated 20 min with the drugs (CNO 20µM, HC-067047 10 µM, in DMSO 0.03% final concentration) before to start the excitability protocol. After-hyperpolarization current (AHP) was assessed in voltage clamp configuration by holding the cell at -60mV with a step of +60mV for 100 ms. TRPV4 currents were assessed by holding the cell at 0mV followed by a 400 ms ramp from −100 mV to +100 mV. The ramp protocol was applied every 5 seconds for 5 minutes (baseline) and then, the TRPV4 inhibitor, HC067047 (10 µM), was applied and cells were recorded for 20 min. Trpv4 current response was obtained by subtracting the current in the presence of HC067047 from the baseline. The synaptic responses were collected with a Multiclamp 700B-amplifier (Axon Instruments, Foster City, CA), filtered at 2.2 kHz, digitized at 5 Hz, and analyzed online using Igor Pro software (Wavemetrics, Lake Oswego, OR).

### RNA extraction, cDNA synthesis, and RT-PCR

Total RNA was extracted using RNeasy Mini Kit (cat 74104) from QIAGEN. The extraction was performed following the details of the kit. After the extraction, RNA quantification was performed using NanoDrop 1000 (Thermo Scientific) and the samples were stored at –80°C until cDNA synthesis. RNA integrity was checked using the Agilent 2100 Bioanalyzer (RIN was always >8). cDNA synthesis for two-step RT-PCR was performed using the QuantiTect Reverse Transcription Kit (cat 205313) from QIAGEN. For each sample, 1 ug of RNA was retrotranscribed in cDNA following the kit instruction. 200 ng of cDNA was used for the RT-PCR analysis using a Sybr Green technology. Plates were processed on the 7900HT systems from Thermo Fisher Scientific, equipped with automated devices for plates loading. (Tecan Freedom EVO). SHANK3 forward primer 5′ acgaagtgcctgcgtctggac 3′, reverse primer 5′ ctcttgccaaccattctcatcagtg 3′; IL-1β forward primer 5’ caaccaacaagtgatattctccatg 3’, reverse primer 5’ gatccacactctccagctgca 3’; TNF-α forward primer 5’ gacgtggaactggcagaagag 3’, reverse primer 5’ gccacaagcaggaatgagaag 3’; TRPV4 forward primer 5’ gtctcgcaagttcaaggact 3’, reverse primer 5’ aaacttacgccacttgtctc 3’; Actin forward primer 5′ agagggaaatcgtgcgtgac 3′, reverse primer 5′ caatagtgatgacctggccgt 3′. Reactions were carried out using iTaq™ Universal SYBR® Green Supermix (Biorad) by 50°C for 2 min, 95°C for 10 min followed by 40 cycles at 95°C for 15 s and 60°C for 1 min. Relative quantification of gene expression was performed according to the ΔΔ-Ct method ^84^.

### FACS sorting and RNA sequencing

Mice were anesthetized in isoflurane and decapitated to dissect fresh brains in ice-cold aCSF (see above). Brains were kept in ice-cold and O_2_ 95%, CO_2_ 5% bubbled aCSF during the preparation of coronal slices, 300um thick using a vibratome. Selected slices were used to manually microdissect the NAc using a total of 4-5 P30 mice for each experiment. The dissected tissue was moved in 1.5ml FACS buffer (L15 added with Glucose 2mg/ml, Bovine Serum Albumin 0,1%, Citrate Phosphate Dextrose 16.7%, DNAseI 10U/ml). After removing the FACS buffer, the tissue was incubated in 400 ul of L15 0.01%Papain (Worthington, #LS003118) and incubated 30’ at +37°C. The tissue was mechanically disrupted pipetting 10 times with a P1000 and a P200 sterile tip, and Papain digestion was blocked adding FACS buffer 0.02% Chicken egg white inhibitor (Sigma, #T9253). The cell suspension was passed through a 70 µm strainer (ClearLine, # 141379C) and spun at 200g for 5’ at +4°C. The precipitate was resuspended in 1ml FACS buffer, this step was repeated a second time to further wash cellular debris. 8 ul of Hoechst (0.1 mg/mL) were added to the sample and incubated for 7’ at +37°C. Before FACsorting we added the 5 ul of the cellular death dye Draq7TM (Viability dye, Far-red DNA intercalating agent, Beckman Coulter, #B25595). The suspension was sorted on an Astrios II cell sorter (Beckam Coulter), enriching for Hoechst stained and Draq7TM non-stained particles. Forward and side scatter were used to exclude smaller cellular debris and duplets. 488nm and 568nm laser excitation were used to separate the desired combinations of cellular population. Each cell population was sorted in FACS buffer and spun down at 200g for 5’ to be dried and snap-frozen in liquid nitrogen before RNA extraction. FACsorting experiments were performed within the Flow cytometry facility at the University of Geneva.

### Sequencing libraries preparation

To prepare cDNA libraries collected frozen tissue was processed using a QIAGEN RNeasy kit (QIAGEN, #74034) to extract RNA and prepare cDNA libraries using SMARTseq v4 kit (Clontech, # 634888) and sequenced using HiSeq 2500 in 100 pairbase length fragments for a minimum of 1 million reads per sample. Sequences were aligned using STAR aligner^85^ using the mouse genome reference (GRCm38). The number of read per transcripts was calculated with the open-source HTseq Python library^86^. All analyses were computed on the Vital-it cluster administered by the Swiss Institute of Bioinformatics. Sequencing experiments were performed within the Genomics Core Facility of the University of Geneva.

### Sequencing analysis

Count tables were normalized to reads per million (RPM) and genes were filtered keeping only those with more than 10 RPM (supplementary information Table S1). DEseq2 package was used to normalize samples to RPM count tables. In **Fig. 3b** we selected differentially expressed genes were selected on a worst-case scenario threshold of 1.5 fold, keeping the data from the replicates corresponding to the pair that gave the minimum fold change between each pair of conditions tested. The full list of results for the worst-case scenario fold change analysis is shown in **Supplementary figure 4a**. We performed PCA analysis in with all of the samples and all of the genes selected above worst-case scenario threshold of 1.5 (855 genes, supplementary information Table S2), these data were normalized by rlog transformation from the DEseq2 package and then used for PCA analysis. SFARI genes (https://gene.sfari.org/tools) belonging to the list of significantly altered genes in AAV-scr*Shank3* versus AAV-sh*Shank3* infected D1R-tom+ and D1R^-^-tom-samples are plotted in **Supplementary figure 4b-c** and have been tested for enrichment using Fisher test in **Fig. 3d**, the 178 worst case scenario differentially expressed genes in AAV-sh*Shank3* versus AAV-scr*Shank3* D1R-tom+ samples, split in sh upregulated and sh downregulated were analyzed for significantly enriched GO:Terms using GOrilla^87^ and the REViGO^88^ online tools, selecting GO:Terms with adjusted P-value lower than 1e-3. Gene expression heatmap in **Fig. 3e** was produced normalizing the rlog transformation of RPM count tables and allowing samples and genes to cluster by Euclidean distance (supplementary information Table S3). All analysis have been made using R, packages used: DEseq2^89^, reshape2^90^, ggplot2^91^, scater^92^, IHW^93^. Count table and FASTQ files are available at the GEO database (GSE139683).

### Cannulations and intra-NAc microinfusions

As explained in the subchapter “Viruses and stereotactic injections”, adult mice (P50-60) were placed on a stereotaxic frame (Angle One; Leica, Germany). Bilateral craniotomy (1 mm in diameter) was then performed bilaterally with the following stereotactic coordinates: AP: +1.2 mm, ML: ± 1 mm, DV: −3.8 mm (measured from bregma). Bilateral stainless steel 26-gauge cannula (5 mm ped, PlasticsOne, Virginia, USA) was implanted above the NAcs and fixed on the skull with dental acrylic. Between experiments, the cannula was protected by a removable cap in aluminum. All animals underwent behavioural experiments 1–2 weeks after surgery. In the rescue experiment with the Trpv4 antagonist, cannulated scr- or sh*Shank3* and *Shank3^+/-^* mice were infused 10 minutes before the three-chamber task (**Fig. 3h**, more precisely, 10 minutes before the habituation in the arena). Cannulated scr- and sh*Shank3* mice performed the behavioural task two times with 7-days pause period between the trails. sh*Shank3* injected mice were randomly infused with 2µL (500nL m^-1^) of vehicle (∼3 % dimethyl sulfoxide (DMSO, Sigma) diluted in aCSF) or with 2µL Trpv4 antagonist (HC-067047, Sigma 2 µg diluted in aCSF-DMSO ∼3 %). The treatment was counterbalanced between trials. On the other hand, scr*Shank3* injected mice were infused both trials with vehicle. Similarly, cannulated *Shank3^+/-^* mice were intraperitoneally injected with LPS at a dose of 2mg/Kg 24 hours before the test. Then mice were randomly infused 10 minutes before the test with 2µL (500nL m^-1^) of vehicle (∼3 % dimethyl sulfoxide (DMSO, Sigma) diluted in aCSF) or with 2µL Trpv4 antagonist (HC-067047, Sigma 2 µg diluted in aCSF-DMSO ∼3 %).

### Tissue processing for post-hoc studies

For *post hoc* analysis, adult mice were anesthetized with pentobarbital (Streuli Pharma) and sacrificed by intracardial perfusion of 0.9% saline followed by 4% PFA (Biochemica). Brains were post-fixed overnight in 4% PFA at 4 °C. 24 hours later, they were washed with PBS before 50μm thick vibratome cutting. After each behavioural experiment, *post hoc* analysis was performed to validate the localization of the infection and/or cannulation.

### Immunohistochemistry and image acquisition

Prepared slices were washed three times with phosphate buffered saline (PBS) 0.1M. Slices were then pre-incubated with PBS-BSA-TX buffer (0.5% BSA and 0.3% Triton X-100) for 90 minutes at room temperature in the dark. Subsequently, cells were incubated with primary antibodies diluted in PBS-BSA-TX (0.5% BSA and 0.3% Triton X-100) overnight at 4°C in the dark. The following day slices were washed three times with PBS 0.1M and incubated for 90 minutes at room temperature in the dark with the secondary antibodies diluted in PBS-BSA buffer (0.5% BSA). Finally, coverslips were mounted using fluoroshield mounting medium with DAPI (Abcam, ab104139). Primary antibody used in this study: polyclonal rabbit anti-Kir3.1 (Girk1, 1/750 dilution, Alamone labs, APC-005). Secondary antibody used at 1/500 dilution: donkey anti-rabbit 488 (Alexa Fluor, Abcam ab150073). *Post hoc* tissue images were acquired using a confocal laser-scanning microscope LSM700 (Zeiss) or an Axiocam fluo wide field microscope (Zeiss) depending on the size of the ROI.

### Statistical analysis

Statistical analysis was conducted with GraphPad Prism 7 and 8 (San Diego, CA, USA) and SPSS version 21.0 (IBM Corp, 2012). Statistical outliers were identified with the ROUT method (Q = 1) and excluded from the analysis. The normality of sample distributions was assessed with the Shapiro–Wilk criterion and when violated non-parametric tests were used. When normally distributed, the data were analyzed with independent t-tests, one sample t-tests, one-way ANOVA and repeated measures (RM) ANOVA as appropriate. When normality was violated, the data were analyzed with Mann–Whitney test, while for multiple comparisons, Kruskal–Wallis or Friedman test was followed by Dunn’s test. For the analysis of variance with two factors (two-way ANOVA, RM two-way ANOVA, and RM two-way ANOVA by both factors), normality of sample distribution was assumed, and followed by Sidak or Tukey post hoc test. Data are represented as the mean ± SEM and the significance was set at 95% of confidence.

